# Pan–Pharmacological Drug–Target Interaction Prediction with 3D–Informed Protein Encoding at Scale

**DOI:** 10.64898/2026.03.27.714727

**Authors:** Akinori Kawaharada, Takumi Ito, Hideyuki Shimizu

**Affiliations:** Department of AI Systems Medicine, M&D Data Science Center, Institute of Integrated Research, Institute of Science Tokyo, Tokyo, JAPAN; Graduate School of Medical and Dental Sciences, Institute of Science Tokyo, Tokyo, JAPAN

**Keywords:** drug–target interaction, binding affinity prediction, multitask learning, protein structure encoding, proteome-wide screening

## Abstract

Accurate prediction of drug–target binding affinity across multiple pharmacological endpoints remains challenging, as most deep learning methodologies focus on a single metric and face a trade–off between incorporating structural information and computational throughput. Here we present OmniBind, a multitask framework that resolves both constraints by encoding protein tertiary structures as discrete token sequences and integrating them with amino acid sequence features through a gated fusion mechanism. Trained on over two million compound–protein pairs from BindingDB, OmniBind simultaneously predicts four pharmacological endpoints in a single forward pass, providing a pan–pharmacological profile across multiple affinity endpoints for each compound–target pair. Across adversarial and temporal benchmarks, OmniBind consistently outperforms state–of–the–art architectures, demonstrating that structural encoding captures physicochemical interaction principles rather than ligand–protein co–occurrence patterns. Attention analysis reveals biologically meaningful binding site recognition, with the model selectively weighting the ABL1 gatekeeper residue T315 and responding to its drug–resistance mutation T315I. As a demonstration of practical utility, proteome–wide screening of 20,421 human proteins recovered 85.7% of known clinical targets of clozapine within the top 200 predictions, correctly resolving its pharmacological profile despite structural similarity to clomipramine. OmniBind provides an accurate, structurally informed, and interpretable platform for multi–endpoint drug-target interaction prediction, with applications to lead optimization, off–target safety assessment, and drug repositioning.

## Introduction

The identification of compounds that modulate a target protein with the desired potency and selectivity lies at the heart of drug discovery. Yet fewer than 10% of candidates successfully navigate the clinical pipeline^1^ despite investments exceeding 2 billion USD per approved therapeutic^2^. A fundamental contributor to this attrition is the combinatorial challenge of navigating a chemical space estimated to exceed 10⁶⁰ molecules^3^ against tens of thousands of human proteins^4^. Accurately predicting binding across the full spectrum of clinically relevant metrics, from fundamental binding affinities (𝐾_𝑖_ and 𝐾_𝑑_) to functional activities (𝐼𝐶_50_ and 𝐸𝐶_50_), is therefore essential for prioritizing compounds at the earliest stages of drug discovery^5^.

Structure–based molecular docking has long served as a rational foundation for this task, but is fundamentally constrained by its requirement for high–resolution experimental structures and its enormous per–pair computational cost, which renders large–scale screening prohibitive^6^. Deep learning has emerged as a high–throughput alternative^7^, yet early architectures based on convolutional or recurrent neural networks were prone to overfitting on small, homogeneous benchmark datasets such as Davis^8^ and KIBA^9^, and failed to generalize to the noise and heterogeneity inherent in real–world experimental data^10^. A more fundamental limitation has persisted across both paradigms: the predominant focus on a single pharmacological metric^11,12,13,14,15^ is insufficient to characterize the multidimensional pharmacological behavior required for clinical relevance.

Achieving such a unified framework requires overcoming three interconnected challenges. First, while AlphaFold2^16^ catalyzed proteome-scale structure prediction, and more recent models such as AlphaFold3^17^ and Boltz–

218 have further advanced structural accuracy, integrating this structural information into deep learning models remains non–trivial. Explicit 3D representations, such as those used by geometric graph networks^19^ or 3D convolutional architectures^20^, dramatically increase computational cost^21^, undermining the high–throughput advantage that motivated deep learning approaches. Second, simultaneous prediction of multiple pharmacological endpoints introduces additional complexity, as differences in data distributions, experimental noise, and measurement principles across these metrics destabilize training and impair generalization^22^. Third, scaling to real–world datasets such as BindingDB^23^, which contains millions of heterogeneous interaction records, demands architectures robust to noise and chemical diversity far beyond that of curated benchmarks. No existing unified framework has resolved all three challenges simultaneously.

Here we present OmniBind, a multitask deep learning framework built around three interlocking design principles that together address this trilemma. First, to incorporate structural information without sacrificing throughput, protein tertiary structure is encoded as discrete one–dimensional 3Di token sequences representing local biophysical environments in a 20–letter structural alphabet^24^, and dynamically integrated with amino acid sequence features through an adaptive gated fusion mechanism^25^, enabling millisecond–scale inference. Second, to capture multi-endpoint pharmacology, OmniBind simultaneously predicts four pharmacological endpoints (𝐾_𝑖_ , 𝐾_𝑑_ , 𝐼𝐶_50_ , and 𝐸𝐶_50_ ) in a single forward pass, providing a pan–pharmacological profile for each compound– target pair. Third, to ensure real–world robustness, the model is trained on over two million interactions from BindingDB^23^, spanning diverse chemical and protein spaces. We show that OmniBind consistently outperforms state–of–the– art models under rigorous adversarial and temporal benchmarks. Attention analysis further reveals that the model learns biologically meaningful binding site representation, and proteome–wide screening demonstrates its utility for off– target profiling of clinical drugs. Together, these capabilities establish OmniBind as a robust, interpretable, and scalable platform for multi-endpoint drug-target interaction prediction.

## Results

### OmniBind architecture: dual–modality protein encoding and multitask drug-target interaction prediction

OmniBind integrates chemical features of compounds with structural and sequential features of proteins through four modules to simultaneously predict multiple pharmacological metrics (**Figure 1**). On the compound side, a graph convolutional neural network extracts atom–level features from molecular graphs from SMILES representations^26,27^. On the protein side, two parallel Transformer encoders independently process amino acid sequences and 3Di sequences generated by ProstT5^28^, capturing sequence–based chemical features and structure–based biophysical features, respectively. The outputs of these two encoders are integrated by a gate fusion layer^25^ (see Materials and Methods) that applies learnable sigmoid gates to dynamically weight the contribution of each modality depending on the binding context. The resulting fused protein representation is combined with the compound representation and processed by a five–layer Transformer decoder, which employs cross–attention to model compound–protein interactions^29^ and outputs predicted values for four pharmacological metrics (𝑝𝐾_𝑖_ , 𝑝𝐾_𝑑_ , 𝑝𝐼𝐶_50_, 𝑝𝐸𝐶_50_) in a parallel manner, which ranges from fundamental binding affinities (𝐾_𝑖_, 𝐾_𝑑_) to cellular efficacy (𝐼𝐶_50_, 𝐸𝐶_50_). To validate the contribution of 3Di integration and the gated fusion mechanism, we performed an ablation study comparing four protein representation strategies (**Table 1**). Single–modality models using only 3Di structural tokens or amino acid sequences achieved comparable performance, indicating that both modalities individually capture informative features for drug-target interaction prediction. Combining the two modalities by simple element– wise addition provided no consistent improvement over either single–modality baseline, suggesting that equal and static weighting of sequence and structural features is insufficient to exploit their complementarity. In contrast, gated fusion achieved the best performance across all four metrics (RMSE, C–index, AUROC, AUPRC), yielding the lowest error (RMSE = 1.037 ± 0.009) and the highest classification accuracy (AUROC = 0.808 ± 0.003, AUPRC = 0.804 ± 0.001). The superiority of gated fusion over simple addition confirms that the two modalities encode complementary rather than redundant information, and that dynamic, context–dependent integration is essential for capturing the distinct biophysical signals encoded in sequence and structural features.

**Table 1:**
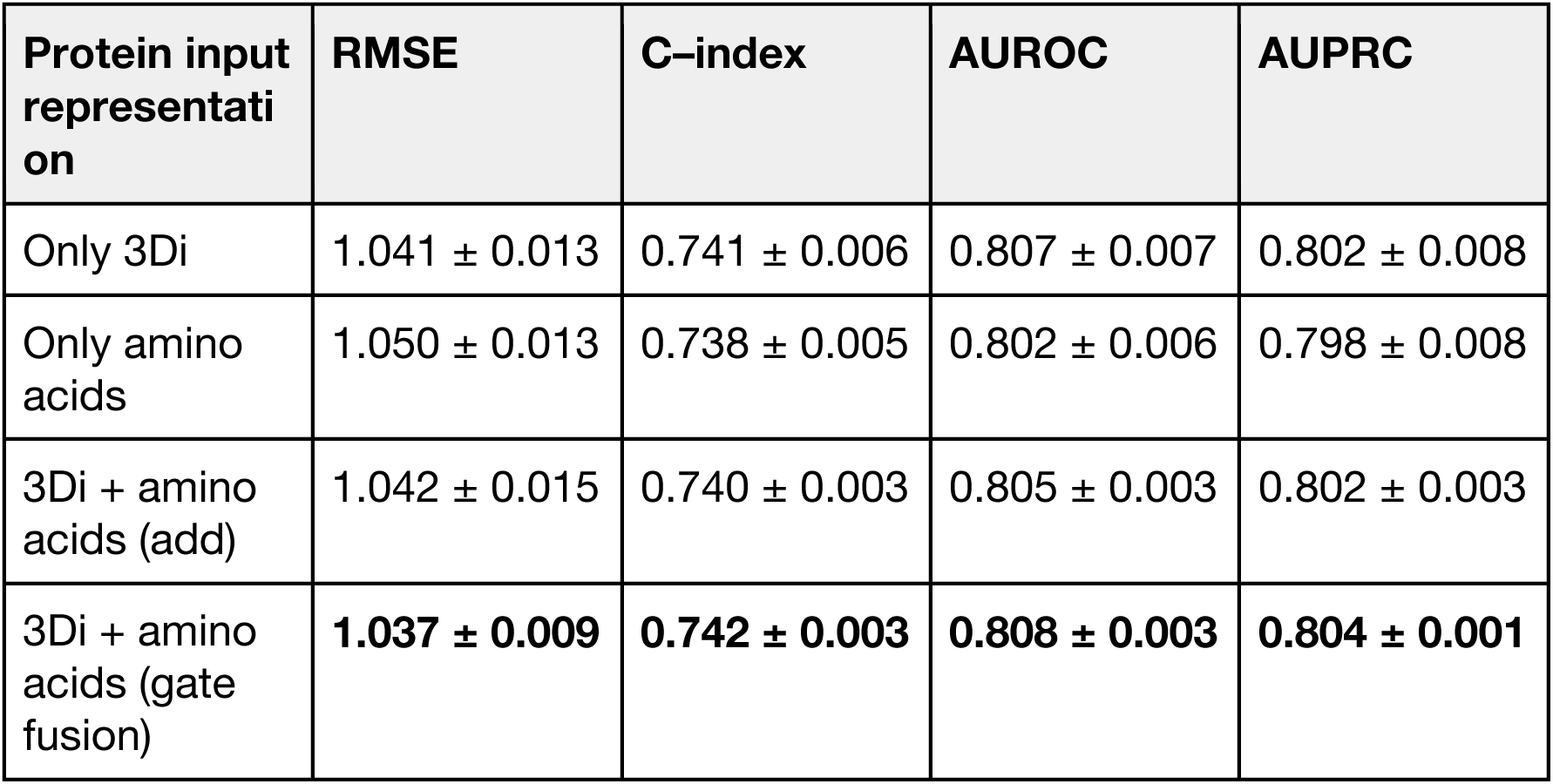
Effect of protein input representation and fusion strategy on drug-target interaction prediction performance. Performance is reported as mean ± standard deviation across five independent runs with different random seeds. "Only 3Di" and "Only amino acids" represent single–modality baselines. "3Di + amino acids (add)" denotes the element–wise addition of the two representations. The "3Di + amino acids (gate fusion)" denotes dynamic integration via learnable sigmoid gates. RMSE, root mean square error; C–index, concordance index; AUROC, area under the receiver operating characteristic curve; AUPRC, area under the precision–recall curve.

**Figure 1:**
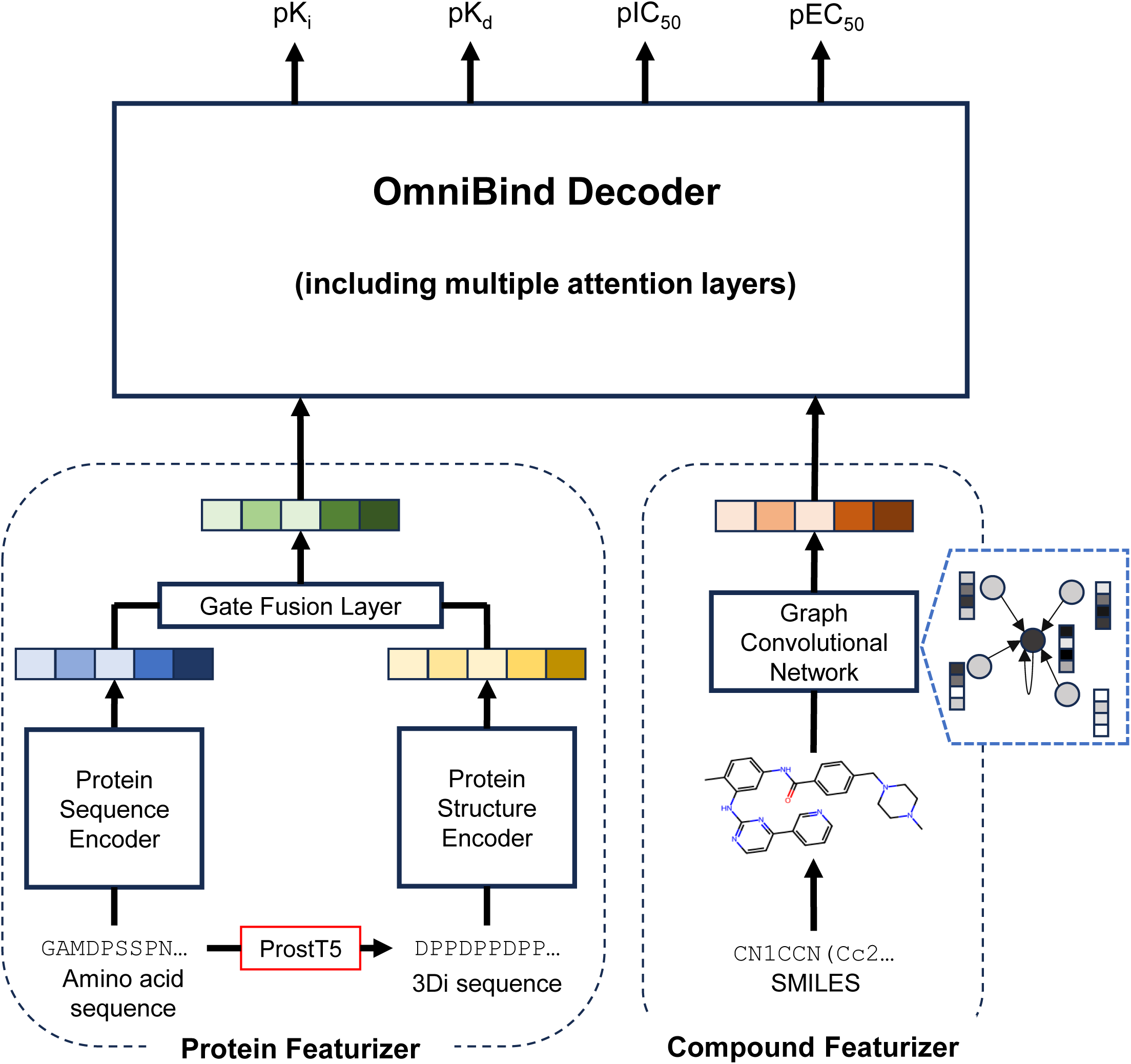
Architecture of OmniBind for pan–pharmacological drug–target interaction prediction. OmniBind processes multimodal protein features and compound topology in parallel before integrating them through a cross–attention decoder. The Protein Featurizer encodes amino acid sequences via Protein Sequence Encoder and 3Di structural token sequences via a Protein Structure Encoder, both implemented as two–layer Transformers. The resulting representations are merged by a gate fusion layer, which applies learnable sigmoid gates to dynamically weight the contribution of each modality rather than concatenating them directly (**Left**). The Compound Featurizer encodes SMILES representations as molecular graphs using a Graph Convolutional Network, producing atom– level feature vectors. The fused protein representation and compound embeddings are then passed to the OmniBind Decoder, a five–layer Transformer that employs cross–attention to model compound–protein interactions and outputs four pharmacological profiles ( 𝑝𝐾_𝑖_, 𝑝𝐾_𝑑_, 𝑝𝐼𝐶_50_ , and 𝑝𝐸𝐶_50_) simultaneously in a single forward pass (**Right**).

### Benchmarking of OmniBind under adversarial and temporal evaluation protocols

To evaluate the generalization capacity of OmniBind on unseen data, we benchmarked it against state–of–the–art models^14,15^ using two complementary evaluation protocols: the label reversal test^13^, which probes whether the model has learned protein–specific interaction features rather than ligand identity, and temporal validation^14^, which assesses generalization to entirely novel chemical and protein spaces by evaluating the model on interactions recorded after the training period, thereby simulating a prospective drug discovery scenario.

In the label reversal test, activity labels are deliberately inverted between training and test sets based on the protein to which each ligand binds (**Figure 2A**), such that any model relying primarily on ligand structure will systematically predict the wrong label^13^. OmniBind achieved a mean RMSE of 1.252 ± 0.032 across the four pharmacological metrics, outperforming DTI–LM^15^ (RMSE = 1.400 ± 0.044) and TransformerCPI2.0^14^ (RMSE = 1.403 ± 0.019) (**Figure 2B**).

**Figure 2:**
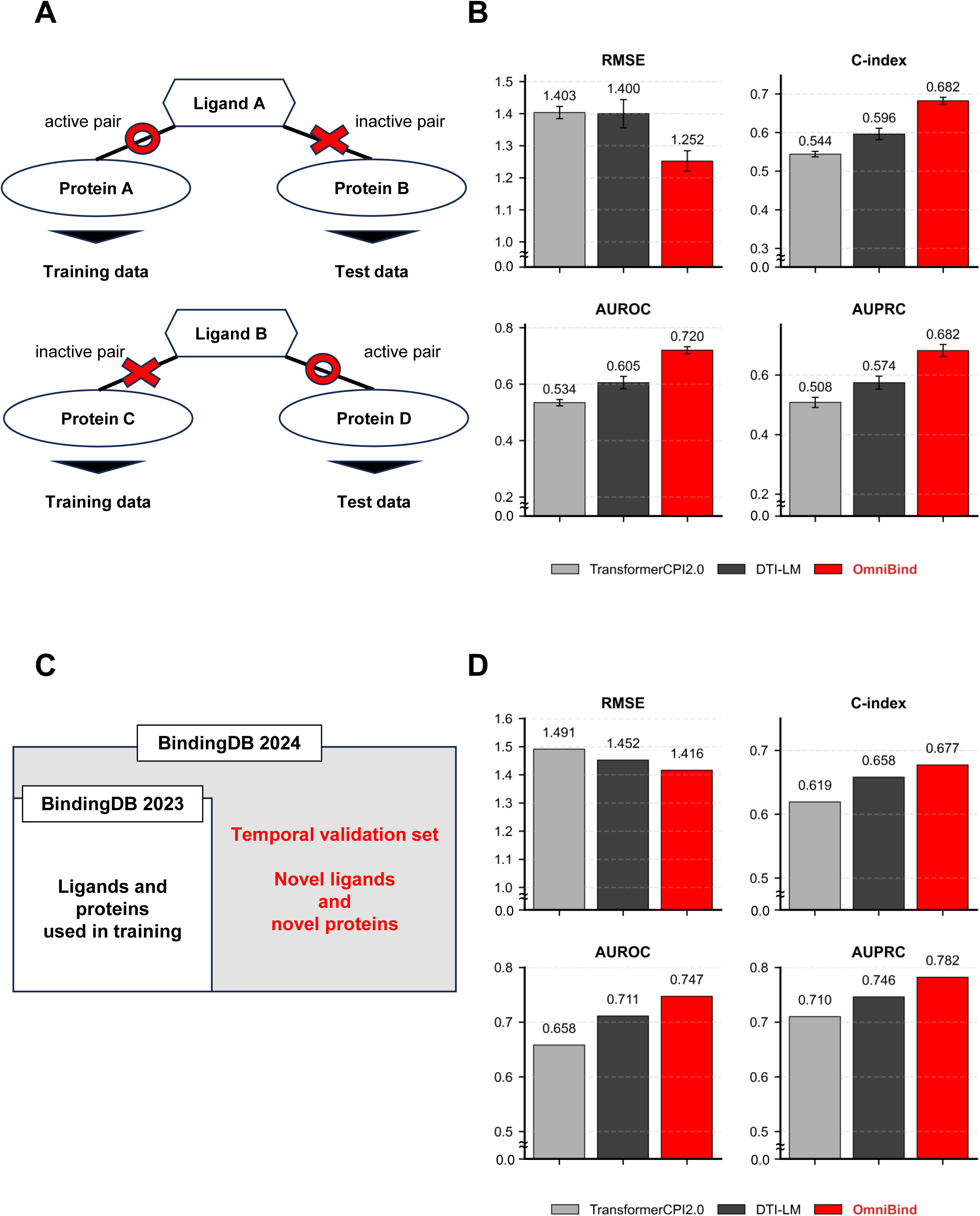
Benchmarking of OmniBind under adversarial and temporal evaluation protocols. **(A)** Schematic of the label reversal test design^13^. Compounds (hexagons) and protein targets (ovals) are partitioned such that the activity label of each ligand is deliberately inverted between training and test sets. For Ligand A (upper panel), only active pairs are included in the training set, while only inactive pairs against different targets are assigned to the test set. For Ligand B (lower panel), the assignment is reversed. Red circles indicate active interactions and red crosses indicate inactive interactions. Under this design, a model relying on ligand structure alone will systematically predict the wrong label, requiring genuine recognition of protein–side features. **(B)** Model performance in the label reversal test. OmniBind (red) outperforms TransformerCPI2.0^14^ (light gray) and DTI–LM^15^ (dark gray) across all four evaluation metrics: RMSE, C–index, AUROC, and AUPRC. Error bars represent the standard deviation across five independent runs. **(C)** Schematic of the temporal validation protocol^14^. The model is trained on BindingDB data^23^ from the May 2023 snapshot and evaluated on interaction pairs newly added to the November 2024 edition (423,647 pairs), with strict exclusion of any compound or protein present in the training set. Ligands and proteins used in training (white area) and the temporal validation set comprising novel ligands and proteins (gray area) are shown. **(D)** Model performance in the temporal validation. OmniBind (red) outperforms TransformerCPI2.0^14^ (light gray) and DTI-LM^15^ (dark gray) across RMSE, C–index, AUROC, and AUPRC, demonstrating generalization to chronologically novel data.

The degraded performance of these large pre–trained models under adversarial conditions suggests that they rely substantially on statistical correlations between compound classes and protein families rather than on the underlying physicochemical principles of molecular recognition^30^.

In temporal validation, the model was trained on BindingDB data^23^ through May 2023 and evaluated on 423,647 interaction pairs newly added to the November 2024 edition, comprising compounds and proteins entirely absent from the training set (**Figure 2C**). OmniBind achieved a mean RMSE of 1.416, outperforming all comparison models across RMSE, C–index, AUROC, and AUPRC (**Figure 2D**), consistent with the results of the label reversal test.

Together, these results demonstrate that OmniBind learns physicochemically grounded interaction representations rather than memorizing dataset–specific patterns. This translates to robust predictive performance under both adversarial and prospective evaluation conditions relevant to real– world drug discovery.

### Drug repositioning screening of FDA–approved drugs and structural validation of predicted interactions

To explore the potential of FDA–approved drugs for novel therapeutic applications, we screened 1,615 FDA–approved drugs against three enzyme targets with distinct functional and clinical backgrounds: phosphodiesterase 5 (PDE5)^31^, plasma kallikrein (KLKB1)^32^, and sirtuin 3 (SIRT3)^33^. These targets were selected to evaluate OmniBind’s predictive versatility across diverse protein classes, spanning a well–characterized phosphodiesterase, a historically challenging serine protease^32,34^, and a relatively underexplored mitochondrial deacetylase (SIRT3)^33^. Top–ranked candidates from OmniBind screening were subsequently validated through a hierarchical pipeline integrating structure prediction by Boltz–2^18^ and docking simulation using MOE^35^, yielding avanafil^36^ as the most promising candidate for PDE5, glecaprevir^37^ for KLKB1, and valrubicin^38^ for SIRT3 (**Figure 3**).

**Figure 3:**
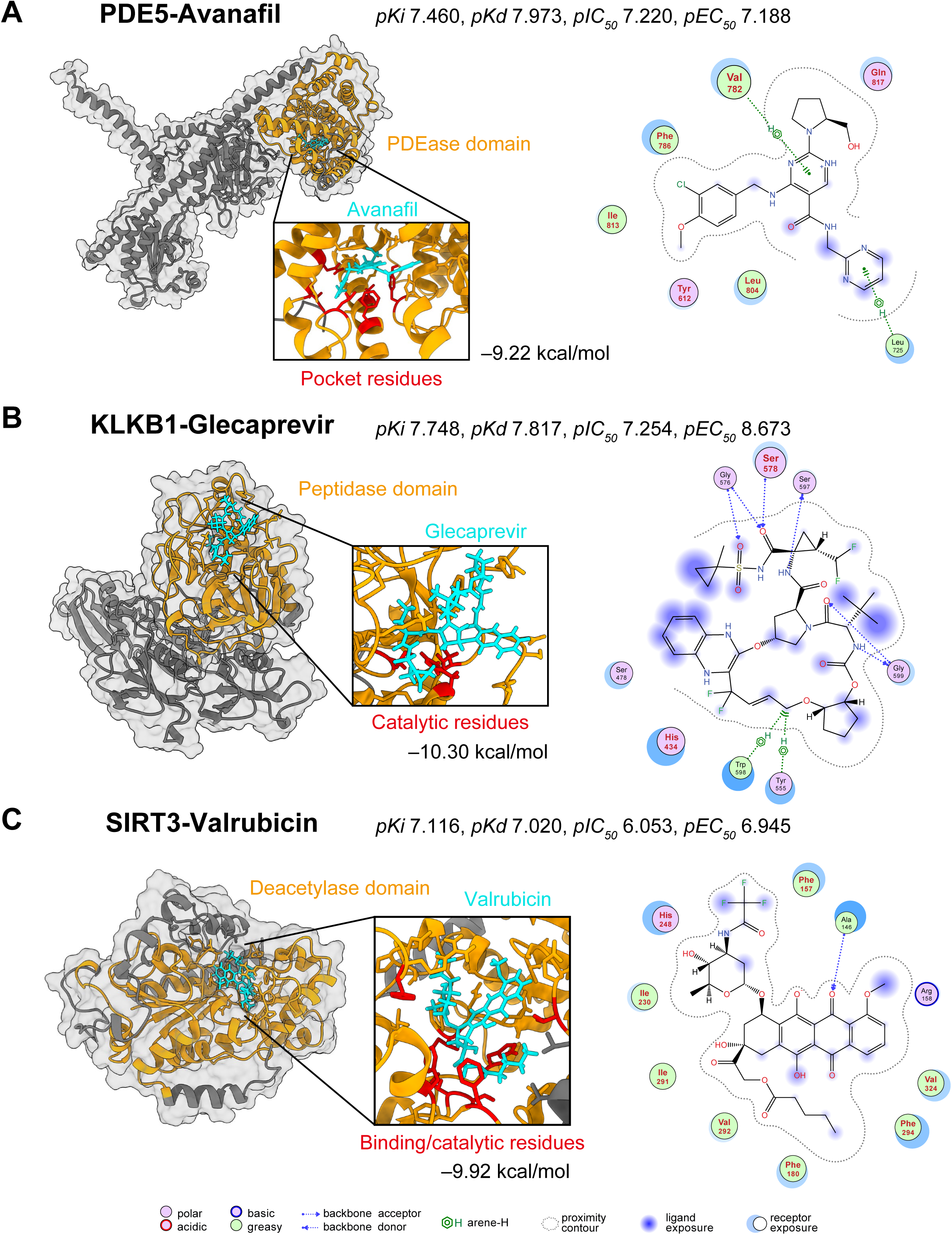
Computational drug repositioning and structural validation of OmniBind–predicted compound–protein interactions. **(A)** Predicted binding of avanafil to PDE5. The compound (cyan) is placed within the PDEase domain (orange), in contact with key binding residues (red) including Val782, a critical residue of the hydrophobic pocket. Avanafil is a known PDE5 inhibitor and was absent from the OmniBind training dataset, serving as an external positive control. The values in the upper right of each panel represent the OmniBind-predicted pharmacological profile, and the value below the 3D structure represents the binding free energy estimated by docking simulation (kcal/mol). The 3D structures of the complexes were visualized using ChimeraX^82^, and the 2D ligand interaction maps were generated using MOE. **(B)** Predicted binding of glecaprevir to KLKB1. The compound (cyan) is positioned within the peptidase domain (orange), in contact with catalytic residues (red) including Ser578, forming a hydrogen bond consistent with serine protease inhibition mechanisms. **(C)** Predicted binding of valrubicin to SIRT3. The compound (cyan) is positioned within the deacetylase domain (orange) such that the acetyl–group binding sites and catalytic residues (red) fall within 4.5 Å of the compound, consistent with the established catalytic mechanism of the sirtuin family.

Avanafil is a known PDE5 inhibitor^36^ and was confirmed to be absent from the OmniBind training dataset, making it an external positive control. Its recovery as the top–ranked candidate for PDE5 demonstrates that OmniBind has learned physicochemically grounded features of the target rather than memorizing training data. Structural modeling places avanafil within the PDEase domain in contact with Val782, a key residue of the hydrophobic pocket, through π– stacking interactions^39^ (**Figure 3A**).

The predicted associations of glecaprevir with KLKB1 and valrubicin with SIRT3 are not currently supported by experimental literature and therefore represent novel drug repositioning candidates. Glecaprevir is positioned within the peptidase domain of KLKB1 in contact with the catalytic residue Ser578^40^, forming a hydrogen bond consistent with serine protease inhibition mechanisms^41^ (**Figure 3B**). Valrubicin is positioned within the deacetylase domain of SIRT3 such that the acetyl–group binding site and catalytic site fall within 4.5 Å of the compound^42^ (**Figure 3C**). Across all three complexes, the predicted binding modes are consistent with the established functional mechanisms of each enzyme^40,42,43^, as corroborated by both Boltz–2 structure prediction and docking simulation.

### Attention analysis reveals biologically meaningful binding site recognition

To assess the interpretability of OmniBind, we analyzed cross–attention weights within the OmniBind Decoder using the ABL1–imatinib interaction as a case study^44,45^ (**Figure 3**). Attention weights in the fourth decoder layer showed a pronounced concentration at the gatekeeper residue T315^46^ (residue 93 in the isolated kinase domain, see Methods), indicating that the model selectively attends to a residue of known functional and clinical significance^47^ (**Figure 3A**). Across decoder layers, a statistically significant difference in mean attention weights between binding and non–binding residues first emerged in Layer 3 (*p* < 0.05, Welch’s *t*–test^48^), reached its maximum separation in layer 4, and attenuated in layer 5 (**Figure 3B**).

**Figure 4:**
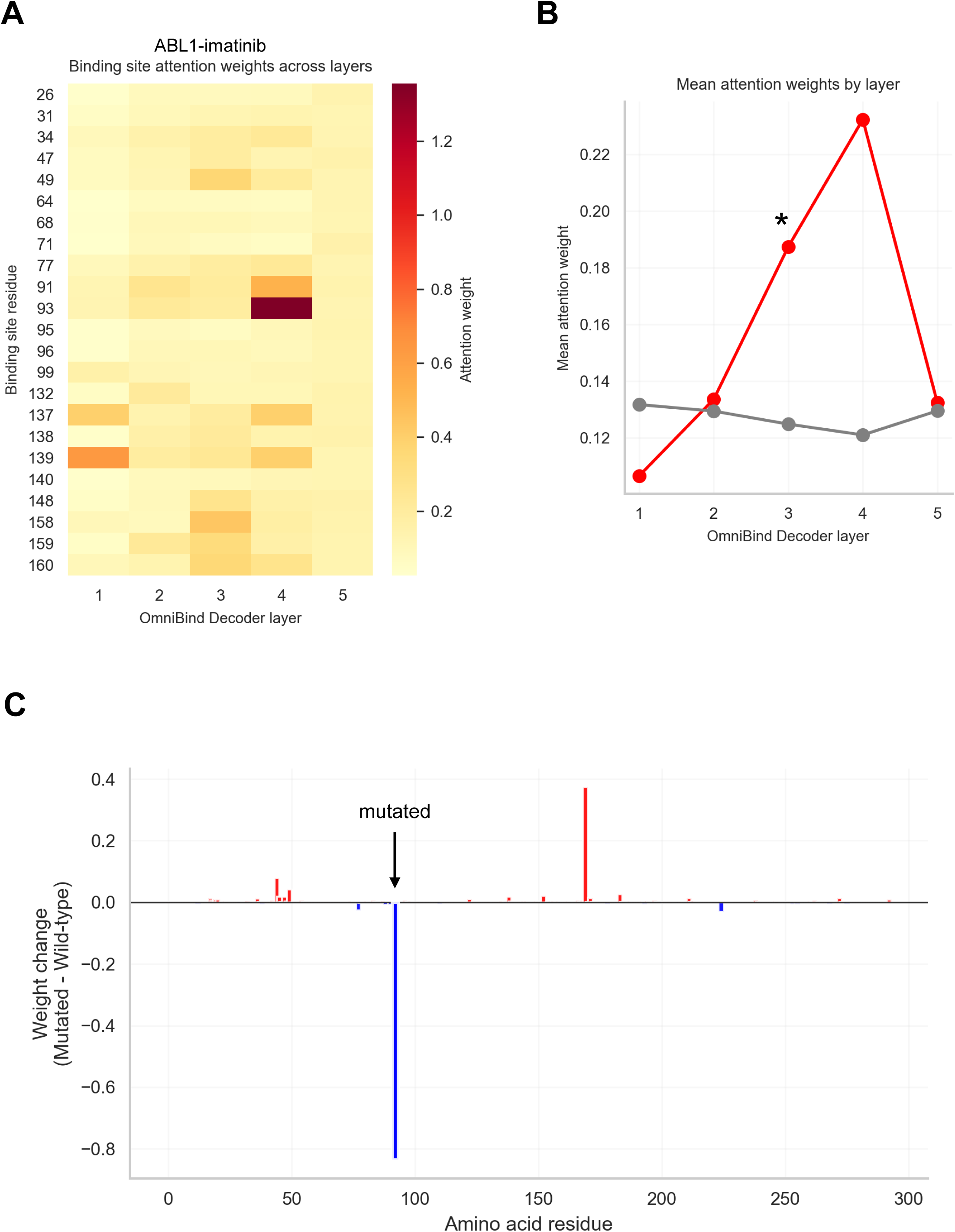
Attention weight analysis of OmniBind decoder layers for the ABL1–imatinib interaction. **(A)** Heatmap of cross-attention weights across the five decoder layers for amino acid residues constituting the ABL1 kinase binding pocket. The gatekeeper residue T315, denoted as position 93 in the isolated kinase domain sequence (see Methods), shows concentrated attention in decoder layer 4. Residue positions correspond to the numbering of the isolated kinase domain (PDB: 6HD6); the full-length ABL1 residue numbering is offset by 222 (e.g., position 93 = T315). **(B)** Mean attention weights assigned to binding site residues (red) and non–binding residues (gray) across each decoder layer, shown for the ABL1– imatinib interaction. A statistically significant difference between the two groups is first observed in Layer 3 (*p* < 0.05, Welch’s *t*–test), and the separation between mean attention weights of binding and non–binding residues reaches its maximum in layer 4. (**C**) Differential attention weights (ΔAttention = mutant – wild–type) across the ABL1 sequence within decoder layer 4 for the T315I substitution. A localized decrease of −0.8352 is observed at residue 93 (arrow), corresponding to the gatekeeper position T315 in full-length ABL1. Blue and red bars represent decreased and increased attention weights in the mutant relative to wild–type, respectively.

To assess the model’s sensitivity to clinically relevant structural perturbations, we introduced the T315I gatekeeper mutation—the most prevalent resistance mutation in imatinib-treated chronic myeloid leukemia patients^49^—and compared attention weights in layer 4 between the wild–type and mutant sequences. The attention weight at residue 93 decreased by 0.8352 in the T315I mutant relative to wild–type (**Figure 3C**), demonstrating that the model detects the structural consequence of this drug–resistance mutation at single–residue resolution^46^. This sensitivity to single-residue substitutions of clinical relevance suggests that attention weights may serve as interpretable indicators of binding site perturbations, with potential utility in predicting resistance-associated changes in drug binding.

To confirm the specificity of these attention patterns, the non–binding MAP2K1–imatinib pair was used as a negative control^50^. The mean attention scores for MAP2K1–imatinib remained consistently lower than those for the ABL1–imatinib across all decoder layers, with no layer–specific signal concentration (**Supplementary Figure S1A**). In contrast to ABL1–imatinib, which showed a prominent peak in maximum attention weights in the fourth decoder layer, the MAP2K1–imatinib pair showed no comparable localized signal across any layer (**Supplementary Figure S1B**). Together, these results demonstrate that the elevated and layer–specific attention patterns observed for ABL1–imatinib reflect genuine molecular recognition rather than a non–specific property of the decoder architecture.

### Proteome–wide off–target profiling demonstrates practical utility of OmniBind

To demonstrate the practical utility of OmniBind in a clinically relevant context, we performed proteome–wide screening against 20,421 UniProt–reviewed human proteins^51^ for two clinical drugs absent from the training dataset: the antipsychotic clozapine^52^ and the antidepressant clomipramine^53^ (**Figure 4A**).

**Figure 5:**
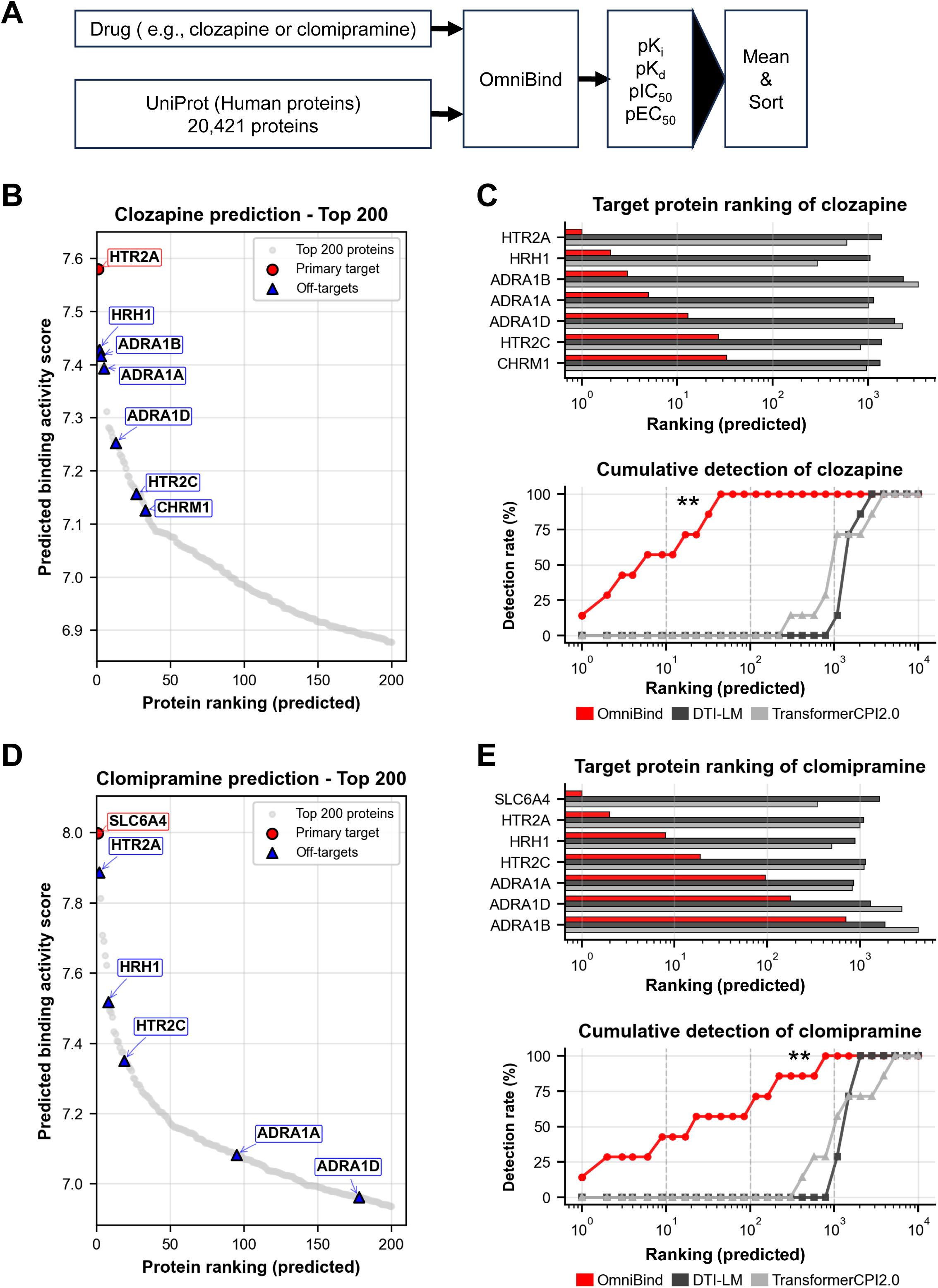
Proteome–wide off–target profiling of clozapine and clomipramine by OmniBind. **(A)** Screening workflow. A query compound and the full UniProt–reviewed human proteome (20,421 proteins) are provided as input. OmniBind simultaneously predicts four pharmacological endpoints ( 𝑝𝐾_𝑖_ , 𝑝𝐾_𝑑_ , 𝑝𝐼𝐶_50_ , 𝑝𝐸𝐶_50_) for each compound–protein pair, and proteins are ranked in descending order of the mean predicted score. (**B**) Predicted target profile of clozapine across the top 200 ranked proteins. As a MARTA with affinity for a broad range of receptor classes, clozapine has multiple known therapeutic and off-targets; HTR2A, which ranked first among all 20,421 proteins, is indicated by a red circle. Known off–targets clinically associated with adverse reactions including sedation and orthostatic hypotension are indicated by blue triangles. (**C**) Comparison of target recovery performance for clozapine among OmniBind (red), DTI–LM^15^ (dark gray) and the TransformerCPI2.0^14^ (light gray). Upper panel: rank distribution of validated targets. Lower panel: cumulative detection rate curves as a function of search space size. OmniBind recovers a greater proportion of known targets within significantly smaller search space (*p* < 0.01 vs. both DTI– LM and TransformerCPI2.0, Wilcoxon signed–rank test). (**D**) Predicted target profile for clomipramine across the human proteome. The primary target SLC6A4 (serotonin transporter)^53^ is ranked first (red circle). Known off–targets including HRH1, HTR2A, HTR2C, and ADRA1A are indicated by blue triangles. (**E**) Comparison of target recovery performance for clomipramine among OmniBind (red), DTI–LM (dark gray), and TransformerCPI2.0 (light gray). Upper panel: rank distribution of validated targets. Lower panel: cumulative detection rate curves. OmniBind recovers a greater proportion of known targets within fewer predictions (*p* < 0.01 vs. both DTI–LM and TransformerCPI2.0, Wilcoxon signed–rank test).

For clozapine, a multi-acting receptor-targeted antipsychotic (MARTA)^54^ known to bind a broad range of receptors^55^, HTR2A (5-hydroxytryptamine receptor 2A) was ranked first among all 20,421 proteins (**Figure 4B**). Known off– targets clinically associated with clozapine–induced adverse effects were also recovered at high ranks, including HRH1 (histamine H1 receptor; rank 4, sedation^56^), ADRA1A and ADRA1B (adrenoceptor alpha 1A/B; ranks 5 and 6, orthostatic hypotension^56^), and CHRM1 (Muscarinic acetylcholine receptor M1; rank 194, anticholinergic effects^56^) ^55^. In total, six of seven known therapeutic and off–targets (85.7%) were recovered within the top 200 predictions, substantially exceeding the detection rates of DTI–LM^15^ and TransformerCPI2.0^14^, neither of which recovered even a single known target within the same search space (**Figure 4C**).

For clomipramine, a serotonin-norepinephrine reuptake inhibitor^53,57^, the primary target SLC6A4 (serotonin transporter)^55^ was ranked first (**Figure 4D**). Known off–targets including HTR2A (rank 2)^56,57^, HRH1 (rank 45)^56^, and ADRA1A (rank 149)^56^were also recovered. Notably, although clozapine and clomipramine share a tricyclic scaffold^55^, OmniBind correctly distinguished their distinct primary targets, HTR2A versus SLC6A4, and reproduced their respective off– target profiles, demonstrating that the model resolves pharmacologically meaningful differences between structurally similar compounds.

These results show that OmniBind can accurately recover clinically validated targets and side–effect–associated off–targets from a proteome–wide search using only sequence–based inputs, supporting its applicability to safety assessment and drug repositioning.

## Discussion

OmniBind addresses three interconnected challenges in deep learning–based drug–target interaction prediction: the generalization to real–world heterogeneous data, the simultaneous prediction of multiple pharmacological metrics, and the efficient integration of protein 3D structural information without sacrificing computational throughput. Incorporating structural information into proteome–wide screening has been particularly difficult, as explicit 3D representations such as geometric graph networks require per–protein computational costs that scale prohibitively when applied to tens of thousands of targets. OmniBind overcomes this barrier by encoding protein tertiary structure as discrete 3Di token sequences^24,28^, enabling structural information to be processed at sequence–level speed and applied to all 20,421 UniProt– reviewed human proteins^51^ in a single screening run. By combining this approach with large–scale training on BindingDB and multitask prediction of four pharmacological endpoints through the gate fusion layer, OmniBind consistently outperformed state–of–the–art models under adversarial and temporal benchmarks, correctly identified known major targets and known off–targets of two clinical drugs in proteome–wide screening, and produced attention patterns that reflect biologically meaningful binding site features. These results establish OmniBind as a robust and practically deployable framework for pan– pharmacological drug–target interaction prediction.

A central technical contribution of OmniBind is the combination of 3Di sequence encoding and adaptive gate fusion for the integration of protein structural and sequence information. Unlike explicit 3D representations such as geometric graph networks, which incur substantial computational cost per protein, 3Di encoding represents protein tertiary structure as a discrete one– dimensional token sequence, allowing structural information to be processed at sequence–level throughput. Importantly, whereas structure-based docking or AlphaFold 3-based interaction prediction typically requires minutes to hours per compound-protein pair, OmniBind completes inference in milliseconds per pair, enabling screening of 20,421 proteins in a single run. Gated fusion further refines this integration by dynamically weighting the relative contribution of structural and sequence features depending on the binding context, rather than combining them by static addition or concatenation. The ablation study confirmed that the two modalities encode distinct yet synergistic features: while amino acid sequences capture global evolutionary signatures, 3Di tokens provide local geometric constraints essential for precise pocket recognition. The gating mechanism’s superiority over simple addition indicates that dynamic context– dependent integration is essential for capturing the complementary information encoded in each representation. This design also positions OmniBind as complementary to high–precision methods such as AlphaFold 3^17^ and structure– based docking: OmniBind is suited for initial high–throughput screening of chemical spaces spanning 10^6^ to 10^8^ compounds, after which top candidates can be subjected to precision structural modeling in a hierarchical workflow^58,59^. The robustness of OmniBind under adversarial benchmarking conditions provides evidence that the model captures features reflecting the physicochemical principles of molecular recognition rather than statistical co– occurrence patterns between compound classes and protein families. This is further supported by attention analysis, which showed that decoder layer 4 selectively weights the ABL1 gatekeeper residue T315. Notably, the model demonstrated single–residue sensitivity by significantly shifting its attention weight in response to the T315I mutation. Such high–resolution responsiveness is particularly valuable given that many global structure–prediction models often struggle to resolve the structural impacts of single–residue substitutions^60^. The ability of internal attention states to detect single–residue structural perturbations of clinical relevance suggests that attention weights may serve as interpretable indicators for binding site features, with potential utility for computationally flagging resistance-associated mutations prior to experimental validation.

The proteome–wide off–target profiling of clozapine and clomipramine illustrates the practical value of OmniBind in a clinically relevant setting. Screening 20,421 human proteins using only sequence–based inputs, OmniBind correctly ranked known major targets of both drugs among the top predictions and recovered 85.7% of known clozapine targets within the top 200 predictions, outperforming DTI–LM^15^ and TransformerCPI2.0^14^. Notably, clozapine is a MARTA with affinity for a broad range of receptor classes^54,55^, and the ability of OmniBind to recover the majority of its known targets while also correctly distinguishing its pharmacological profile from the structurally similar tricyclic compound clomipramine demonstrates compound-level specificity that is directly relevant to safety assessment and drug repositioning.

As with all data–driven models, OmniBind is subject to the quality and coverage biases of BindingDB^23^, which may underrepresent certain protein families or chemical scaffolds. Incorporating more diverse data sources or physicochemical constraints through approaches such as physics–informed neural networks^61^ could improve performance in underrepresented regions of chemical and protein space. Additionally, the current model does not account for conformational dynamics, post–translational modifications, or allosteric effects that influence binding *in vivo*. Integration of dynamic structural information from molecular dynamics simulations represents a potential avenue for improving predictive accuracy. Looking ahead, integration of OmniBind as a scoring function within molecular generative models through reinforcement learning or genetic algorithms could enable closed–loop *de novo* drug design workflows^62^, with attention–based interpretability providing insight into the biophysical basis of predicted interactions.

In conclusion, OmniBind demonstrates that lightweight 3D structural encoding combined with gated fusion and multitask learning enables drug– target interaction prediction that is robust to ligand bias, generalizable to novel chemical and protein spaces, and scalable to proteome–wide applications. By making structural information accessible at high–throughput screening scale, OmniBind provides a practical computational platform for lead optimization, off– target safety assessment, and drug repositioning.

## Materials and Methods

### Dataset Construction

Two editions of BindingDB^23^ were used as the primary data source: May 2023 for model development and November 2024 for temporal validation. BindingDB was selected because it offers the broadest coverage of experimentally measured compound–protein binding interactions among publicly available databases, encompassing diverse assay types, protein families, and chemical scaffolds, conditions that are essential for training a model intended for real– world drug discovery applications^23^. To avoid information leakage resulting from naive random splitting, we constructed two independent test sets with distinct evaluation objectives: a label reversal test set to assess ligand bias^13^, and a temporal validation set for time–based generalization assessment (see below for the details)^14^.

### Data Cleaning

Raw BindingDB underwent the following preprocessing steps. First, only single– chain proteins were retained, as multi–chain complexes introduce structural ambiguity that cannot be adequately represented by sequence–based encoders. Entries were kept only if both a SMILES representation and an amino acid sequence were present, with at least one of the four pharmacological metrics (𝐾_𝑖_, 𝐾_𝑑_, 𝐼𝐶_50_, or 𝐸𝐶_50_) recorded. Inequality signs (e.g., ">" or "<") were removed to convert activity values into numerical form, and outliers exceeding 10^7^ nM were excluded as these likely reflect assay artifacts rather than genuine binding events. All activity values were converted from nM units to 𝑝𝐴𝑐𝑡𝑖𝑣𝑖𝑡𝑦 values (−𝑙𝑜𝑔_10_(𝑀) ) to place all four metrics on a common scale and stabilize regression training^63^. Invalid SMILES representations that could not be parsed into mol objects by RDKit (version 2022.9.5)^64^ were excluded, and duplicate compounds were consolidated into unique canonical SMILES to prevent redundancy in training data. Protein sequences containing hyphens or non–IUPAC characters were also removed, as these indicate modified residues or annotation errors incompatible with standard sequence encoders. After preprocessing, 2,282,997 pairs were retained from the May 2023 edition and 2,761,182 pairs from the November 2024 edition. Dataset statistics for each metric are provided in **Supplementary Table S1**.

### Label Reversal Test Set

To assess whether OmniBind genuinely learns protein–specific interaction features rather than relying on ligand identity alone, a form of overfitting known as ligand bias, we constructed a label reversal test set following the protocol proposed in TransformerCPI study^13^. From the May 2023 BindingDB snapshot, 100,000 unique ligands were extracted that had both active (𝑝𝐴𝑐𝑡𝑖𝑣𝑖𝑡𝑦 ≥ 6.5) and inactive ( 𝑝𝐴𝑐𝑡𝑖𝑣𝑖𝑡𝑦 < 6.5 ) annotations across different proteins. These ligands were randomly divided into two equal groups of 50,000. For the first group, active pairs were assigned to the test set and inactive pairs to the training set, while the assignment was reversed for the second group. Under this adversarial design, the activity label of any given ligand is deliberately inverted between training and test, meaning a model that relies primarily on ligand structure will systematically predict the wrong label. Genuine generalization therefore requires the model to capture protein–side features that distinguish active from inactive binding contexts. Remaining data outside the label reversal partition were split into training and validation sets with no ligand overlap between partitions. Detailed statistics for each partition are provided in **Supplementary Table S1**.

### Temporal Validation Set

To evaluate generalization to compounds and proteins absent from the training data, thereby simulating a prospective drug discovery scenario, we performed a time–based split using the two BindingDB editions^23^. The model was trained on the May 2023 snapshot, and interaction records newly added to the November 2024 edition were used as the test data. To eliminate any possibility of information leakage, test pairs were strictly filtered to retain only those in which both the compound and the protein were absent from the training set, ensuring that neither molecular nor proteomic overlap existed between training and evaluation. This partitioning resulted in a temporal validation set comprising 423,647 compound–protein pairs. Time–based splitting provides a more realistic estimate of predictive performance than random splitting, as it mirrors the prospective nature of real–world drug discovery where models must generalize to interactions not yet characterized at the time of training^14^. Dataset statistics for each activity metric are summarized in **Supplementary Table S1**.

### Data Preprocessing and Feature Engineering

Compounds were obtained as SMILES strings and converted into canonical SMILES using the RDKit library (version 2022.9.5)^64^ to ensure structural uniqueness and prevent duplicate representations across the datasets. To incorporate three–dimensional (3D) structural information without requiring experimental structures, 3Di sequences were generated from amino acid sequences using the protein language model ProstT5^28^. The 3Di alphabet encodes the local structural environment of each residue into one of 20 discrete tokens, enabling lightweight representation of protein tertiary structure at sequence–level computational cost^24^. Although 3Di sequences derived from predicted rather than experimental structures may contain minor errors, consistently utilizing sequence–derived 3Di ensures that OmniBind can be applied to any protein regardless of the availability of experimental structures, substantially broadening its practical applicability. Experimental activity values for 𝐾_𝑖_, 𝐾_𝑑_, 𝐼𝐶_50_ , and 𝐸𝐶_50_ were converted into 𝑝𝐴𝑐𝑡𝑖𝑣𝑖𝑡𝑦 ( 𝑝𝑋 = − log_10_(𝑋) ) to place all metrics on a common logarithmic scale, yielding 𝑝𝐾_𝑖_, 𝑝𝐾_𝑑_, 𝑝𝐼𝐶_50_, and 𝑝𝐸𝐶_50_respectively. For binary classification tasks, continuous 𝑝𝐴𝑐𝑡𝑖𝑣𝑖𝑡𝑦 values were binarized at a threshold of 6.5, following the protocols established in TransformerCPI study^13,14^.

### OmniBind Model Architecture

OmniBind consists of a compound featurizer, a protein featurizer with a gate fusion layer, and a Transformer decoder that integrates both representations to predict multiple pharmacological endpoints simultaneously (**Figure 1**).

### Compound Featurizer

Molecular graphs are encoded using a single–layer Graph Convolutional Network (GCN)^27^, in which each atom is represented by a 35–dimensional feature vector calculated via RDKit (**Supplementary Table S2**). A virtual atom connected to all other atoms in the graph was introduced following the design of TransformerCPI2.0^14^, enabling the aggregation of global molecular context alongside local atom–level features. The number of GCN layers was fixed at one, as preliminary experiments consistent with the findings of TransformerCPI2.0^14^ showed that additional layers cause over–smoothing of atomic features, degrading predictive performance.

### Protein Featurizer

Amino acid sequences and 3Di structural token sequences are encoded by two independent two–layer Transformer encoders, the Protein Sequence Encoder and the Protein Structure Encoder, respectively. Each encoder layer uses 8– head multi–head self–attention with a hidden dimension of 256 and a feed–forward dimension of 1024. GELU activation^65^ and Layer Normalization^66^ were applied throughout to stabilize training.

The output [CLS] tokens from both encoders, denoted as h_pc_ (sequence) and h_ps_ (structure), are passed to the gate fusion layer. Rather than merging the two representations by simple addition or concatenation, which treats both modalities equally regardless of the binding context, the gate fusion layer computes learnable sigmoid gate weights (𝑝_𝑝𝑐_, 𝑝_𝑝𝑠_) to dynamically modulate the relative contribution of each modality. The weights and biases of the linear layers within the fusion mechanism were initialized using the PyTorch (version 2.1.0)^67^ default uniform distribution:

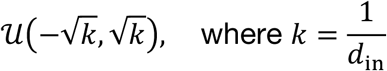

The fusion is defined by the following equations:

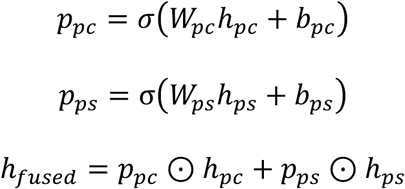

where 𝑊_𝑝𝑐_, 𝑊_𝑝𝑠_ ∈ ℝ^𝑑×𝑑^ and 𝑏_𝑝𝑐_, 𝑏_𝑝𝑠_ ∈ ℝ^𝑑^ are learnable weight matrices and bias vectors, σ denotes the sigmoid activation function, and ⊙ represents the Hadamard product (**Supplementary Figure S2A**).

### OmniBind Decoder

A five–layer Transformer decoder receives the atom–level compound embeddings from the compound featurizer and the fused protein representation from the gate fusion layer. Within each decoder layer, multi-head self-attention is first applied to the compound embeddings, followed by multi-head cross–attention between the compound embeddings and the fused protein representation, allowing the decoder to model compound–protein interactions in a context–dependent manner. The vector corresponding to the virtual atom is then passed through a fully connected layer to simultaneously output predicted values for all four pharmacological endpoints. The decoder shares the same architectural hyperparameters as the encoders (8 attention heads, hidden dimension of 256). The full decoder architecture is illustrated in **Supplementary Figure S2B**.

### Hyperparameter Tuning

Hyperparameters were determined through a stepwise grid search^68^. The numbers of encoder and decoder layers were optimized first, followed by the hidden dimension and attention head count, and finally dropout rate^69^. This sequential strategy was adopted to reduce the combinatorial search space given computational resource constraints. To accelerate each search phase, a 25% random subsample of the training and validation datasets was used, with validation RMSE as the selection criterion. The final hyperparameter configuration is summarized in **Supplementary Table S3**.

### Training and Evaluation

Model training was performed using the RAdam optimizer^70^ with a learning rate of 1 × 10^−4^ and a weight decay of 1 × 10^−4^. RAdam was selected over standard Adam because it rectifies the variance of the adaptive learning rate in early training stages, providing more stable convergence on large, heterogeneous datasets. Due to the memory demands of processing full protein sequences through Transformer encoders, a per–device batch size of 2 was used with gradient accumulation over 64 steps, resulting in an effective batch size of 128.

Because BindingDB entries do not uniformly report all four pharmacological metrics, a masking strategy was applied to the loss function to handle missing values. For each batch, the mean squared error (MSE) loss was calculated independently for each available endpoint and averaged over the number of metrics present in that batch. For instance, if a batch contains only 𝐾_𝑖_, 𝐼𝐶_50_, and 𝐸𝐶_50_ measurements, the loss was defined as the mean of the three corresponding MSE values. No manual weighting was applied across metrics, as differential weighting risks over–optimizing toward metrics with larger sample sizes and may impair the generality of the learned representations.

Distributed training was conducted using the Horovod library^71^ across eight NVIDIA A100 GPUs (40 GB each), completing 100 epochs in approximately 60 hours. Convergence was confirmed by the steady decline in both training and validation RMSE and a concurrent increase in the validation C–index over the full training period (**Supplementary Figure S3**). Regression performance was evaluated using RMSE and the C–index^72^, while classification performance was assessed using AUROC and AUPRC^73^. Primary performance metrics reported in **Table 1** represent the mean across three primary metrics: 𝑝𝐾_𝑖_, 𝑝𝐼𝐶_50_ and 𝑝𝐸𝐶_50_. Although 𝑝𝐾_𝑑_ was predicted simultaneously, it was excluded from the primary average due to its substantially smaller sample size and higher inter–replicate variance in BindingDB would destabilize cross–model comparisons. Individual performance for all four labels including 𝑝𝐾_𝑑_is reported in **Supplementary Table S4** for completeness.

To ensure a fair comparison across existing models^15,14^, all methods were retrained and evaluated on a common dataset. Entries with SMILES strings exceeding 500 characters or amino acid sequences exceeding 1500 residues were excluded to accommodate the memory constraints of DTI–LM^15^. This filtering removed 4.5% of BindingDB entries. To confirm that this exclusion did not affect the validity of our conclusions, OmniBind was separately evaluated on the excluded long–sequence entries alone, and no statistically significant difference in mean RMSE was observed compared to the performance on the full dataset (Wilcoxon signed–rank test^74^, *p* = 0.438).

### Drug Repositioning and Structural Validation

#### Target Selection and Initial Screening

To explore the potential of FDA–approved drugs for novel therapeutic indications, we predicted the binding affinities against three enzyme targets with distinct structural, functional, and clinical backgrounds: phosphodiesterase 5 (PDE5)^31^, plasma kallikrein (KLKB1)^32^, and sirtuin 3 (SIRT3)^33^. PDE5 is a well–established target for erectile dysfunction whose isoform selectivity is critical for minimizing side effects, making it a useful positive control for evaluating whether OmniBind captures essential structural features and binding principles of a known catalytic pocket. KLKB1 is a serine protease involved in blood pressure regulation and inflammatory pathways and represents a protein class that is historically challenging for small–molecule drug development^75^. SIRT3 is a mitochondrial NAD⁺–dependent deacetylase essential for cellular metabolism and stress response, and was included as a less–characterized therapeutic target to assess the model’s ability to generalize beyond well–studied protein families^76^. Together, these three targets span distinct enzyme classes and provide a stringent test of OmniBind’s versatility across diverse compound–protein interaction landscapes. For each target, the top 30 compounds ranked by their mean OmniBind scores (𝑝𝐾_𝑖_, 𝑝𝐼𝐶_50_, 𝑝𝐸𝐶_50_) were selected for high–precision structural modeling.

#### Protein–Ligand Structure Prediction via Boltz–2

Three–dimensional complex structures of the top–ranked candidates were predicted using Boltz–2^18^. Boltz–2 was selected because it provides rapid and accurate prediction of protein–ligand binding poses and estimates binding affinity probability, enabling an efficient filtering step prior to detailed docking.

Predictions were run without structural templates to avoid bias toward known binding modes, utilizing Multiple Sequence Alignment (MSA) input and potential– based scoring to ensure conformational diversity. Diffusion sampling steps and recycling iterations were set to five each, following the default parameters recommended for ligand binding prediction. From the resulting structural ensembles, poses with an affinity probability of ≥ 0.5 were retained for downstream docking analysis, as this threshold corresponds to complexes predicted to be genuine binders with greater than chance confidence.

#### Molecular Docking and Refinement in MOE

Retained complexes were visualized in the Molecular Operating Environment (MOE, Chemical Computing Group Inc.)^35^ to assess compound orientation relative to known functional domains and active sites annotated in UniProt. This visual inspection step was included to exclude poses in which compounds were positioned on solvent–exposed surfaces outside established active domains, which are unlikely to represent pharmacologically relevant binding modes. Compounds satisfying this criterion were carried forward to docking simulation. Structure preprocessing was performed using the QuickPrep module, which applies structural correction, protonation and charge neutralization, and energy minimization of the binding site. Docking was executed using the Triangle Matcher placement algorithm, generating 50 initial poses per compound. Poses were ranked using the London dG scoring function, and the top 10 were retained for refinement under a Rigid Receptor configuration with ligand flexibility. The final binding affinity was estimated using the GBVI/WSA dG scoring function^77^, which approximates the binding free energy by accounting for generalized Born solvation and weighted surface area contributions.

### Attention Analysis

To assess whether OmniBind captures biologically meaningful interaction features, we analyzed the cross-attention weights of the OmniBind decoder using the ABL1 kinase (PDB ID: 6HD6)^78^ and its selective inhibitor imatinib (ChEMBL ID: 941)^79^ as a case study^44,45^. ABL1 was chosen for three reasons: it is a clinically important target in chronic myeloid leukemia with several FDA– approved inhibitors^80^, its high–resolution crystal structures provide well–defined binding site annotations^47^, and the T315I gatekeeper mutation responsible for acquired imatinib resistance is mechanistically well–characterized^46^, providing a stringent test of whether the model responds appropriately to a clinically relevant structural perturbation^41^.

The amino acid sequences were retrieved from the PDB, and the 3Di sequences were generated by submitting the amino acid sequences to the Foldseek server^24^. Because 6HD6 represents the isolated kinase domain in which residues 7 to 293 correspond to residues 229 to 515 of the full–length ABL1, the T315I mutant sequence was constructed by substituting threonine with isoleucine at position 93 of the 6HD6 sequence, equivalent to position 315 in the full–length protein. The SMILES representation for imatinib was obtained from ChEMBL^79^. The amino acid sequences, 3Di sequences, and SMILES string were input into OmniBind to retrieve the predicted pharmacological values and cross-attention weight matrices from all five decoder layers.

Cross-attention weights were obtained as two–dimensional arrays of shape (number of compound atoms) × (number of protein residues). To enable comparison of attention patterns across decoder layers and to quantify changes induced by the T315I mutation, the compound–side dimension was summed and normalized to produce a one–dimensional residue–level attention profile for each layer.

### Off–target Screening

Proteome–wide off–target screening was performed against all UniProt– reviewed human proteins (20,421 proteins)^51^ using clozapine and clomipramine as query compounds, both of which were confirmed to be absent from the training dataset. These two drugs were selected because, despite sharing a tricyclic scaffold^55^, they exhibit distinct primary mechanisms of action^52,53^, providing a stringent test of whether OmniBind can distinguish subtle structural differences and correctly resolve compound–specific pharmacological signatures. For each protein, amino acid sequences were converted into 3Di sequences using ProstT5^28^, and both sequence types were provided as inputs to OmniBind. Because OmniBind simultaneously predicts four pharmacological indicators (𝑝𝐾_𝑖_, 𝑝𝐾_𝑑_, 𝑝𝐼𝐶_50_, and 𝑝𝐸𝐶_50_), the mean of these four predicted values was used as a composite binding score, and proteins were ranked in the descending order of this score. This composite score was adopted as a practical proxy for binding propensity, enabling proteome-wide ranking across all four endpoints simultaneously.

For benchmarking, DTI–LM^15^ and TransformerCPI2.0^14^ were selected as the primary comparison models, consistent with the performance evaluations conducted in previous sections. As DTI–LM and TransformerCPI2.0 are designed for single-endpoint prediction, ranking was performed using 𝑝𝐼𝐶_50_values alone, which is the most commonly reported metric in BindingDB and drug discovery literature. Both models were evaluated based on their ability to recover known therapeutic targets and side–effect–related off–targets within the top–ranked predictions.

Known targets were defined on the basis of PubChem BioAssay data^81^ and published pharmacological literature. For clozapine, a MARTA with affinity for a broad range of receptor classes^54^, the experimentally validated targets comprised HTR2A, HRH1, ADRA1A, ADRA1B, CHRM1, and HTR2C^55,56^. For clomipramine, a serotonin-norepinephrine reuptake inhibitor^53^, known targets comprised SLC6A4 (primary target), HTR2A, HRH1, HTR2C, and ADRA1A, and ADRA1D^55,57^. Detection performance was assessed by comparing the ranks assigned to these known targets by OmniBind, DTI–LM and TransformerCPI2.0.

### Experimental Environment and Reproducibility

All experiments in this study were implemented in Python (ver. 3.8.12) using PyTorch (ver. 2.1.0)^67^ and RDKit (ver. 2022.9.5)^64^. To ensure the reproducibility, five different random seeds (42, 123, 369, 777, and 2024) were employed for both dataset splitting and model weight initialization, and results are reported as mean ± standard deviation across five independent runs.

## ACKNOWLEDGEMENTS

This work was supported by KAKENHI grants from the Japan Society for the Promotion of Science (JSPS) to H.S. (23K18502, 24H01755 and 25H01571), JST FOREST Program to H.S. (JPMJFR242Q), as well as Takeda Science Foundation and Nakatani Foundation to H.S. We thank T. Suzuki, T. Sakuma, Y. Otani, and T. Fujiwara for critical reading of the manuscript, all the laboratory members for discussions, and K. Tanaka for help with preparation of the manuscript.

## AUTHOR CONTRIBUTIONS

H.S. conceived of, designed, and supervised the study. A.K. developed OmniBind and performed computational analyses with the help of T.I., who assisted with 3D molecular modeling. A.K and H.S. jointly wrote the manuscript.

## COMPETING INTERESTS

The authors declare no competing interests.

## Notes

### Competing Interest Statement

The authors have declared no competing interest.

## References

(1) Sun, D.; Gao, W.; Hu, H.; Zhou, S. Why 90% of Clinical Drug Development Fails and How to Improve It? Acta Pharm. Sin. B 2022, 12 (7), 3049–3062. 10.1016/j.apsb.2022.02.002.

(2) DiMasi, J. A.; Grabowski, H. G.; Hansen, R. W. Innovation in the Pharmaceutical Industry: New Estimates of R&D Costs. J. Health Econ. 2016, 47, 20–33. 10.1016/j.jhealeco.2016.01.012.

(3) Bohacek, R. S.; McMartin, C.; Guida, W. C. The Art and Practice of Structure-Based Drug Design: A Molecular Modeling Perspective. Med. Res. Rev. 1996, 16 (1), 3–50. 10.1002/(SICI)1098-1128(199601)16:1%3C3::AID-MED1%3E3.0.CO;2-6.

(4) Ruddigkeit, L.; van Deursen, R.; Blum, L. C.; Reymond, J.-L. Enumeration of 166 Billion Organic Small Molecules in the Chemical Universe Database GDB-17. J. Chem. Inf. Model. 2012, 52 (11), 2864–2875. 10.1021/ci300415d.

(5) Hopkins, A. L. Network Pharmacology: The next Paradigm in Drug Discovery. Nat. Chem. Biol. 2008, 4 (11), 682–690. 10.1038/nchembio.118.

(6) Pinzi, L.; Rastelli, G. Molecular Docking: Shifting Paradigms in Drug Discovery. Int. J. Mol. Sci. 2019, 20 (18), 4331. 10.3390/ijms20184331.

(7) Wen, M.; Zhang, Z.; Niu, S.; Sha, H.; Yang, R.; Yun, Y.; Lu, H. Deep-Learning-Based Drug–Target Interaction Prediction. J. Proteome Res. 2017, 16 (4), 1401–1409. 10.1021/acs.jproteome.6b00618.

(8) Davis, M. I.; Hunt, J. P.; Herrgard, S.; Ciceri, P.; Wodicka, L. M.; Pallares, G.; Hocker, M.; Treiber, D. K.; Zarrinkar, P. P. Comprehensive Analysis of Kinase Inhibitor Selectivity. Nat. Biotechnol. 2011, 29 (11), 1046–1051. 10.1038/nbt.1990.

(9) Tang, J.; Szwajda, A.; Shakyawar, S.; Xu, T.; Hintsanen, P.; Wennerberg, K.; Aittokallio, T. Making Sense of Large-Scale Kinase Inhibitor Bioactivity Data Sets: A Comparative and Integrative Analysis. J. Chem. Inf. Model. 2014, 54 (3), 735–743. 10.1021/ci400709d.

(10) Sieg, J.; Flachsenberg, F.; Rarey, M. In Need of Bias Control: Evaluating Chemical Data for Machine Learning in Structure-Based Virtual Screening. J. Chem. Inf. Model. 2019, 59 (3), 947–961. 10.1021/acs.jcim.8b00712.

(11) Öztürk, H.; Özgür, A.; Ozkirimli, E. DeepDTA: Deep Drug–Target Binding Affinity Prediction. Bioinformatics 2018, 34 (17), i821–i829. 10.1093/bioinformatics/bty593.

(12) Nguyen, T.; Le, H.; Quinn, T. P.; Nguyen, T.; Le, T. D.; Venkatesh, S. GraphDTA: Predicting Drug–Target Binding Affinity with Graph Neural Networks. Bioinformatics 2021, 37 (8), 1140–1147. 10.1093/bioinformatics/btaa921.

(13) Chen, L.; Tan, X.; Wang, D.; Zhong, F.; Liu, X.; Yang, T.; Luo, X.; Chen, K.; Jiang, H.; Zheng, M. TransformerCPI: Improving Compound–Protein Interaction Prediction by Sequence-Based Deep Learning with Self-Attention Mechanism and Label Reversal Experiments. Bioinformatics 2020, 36 (16), 4406–4414. 10.1093/bioinformatics/btaa524.

(14) Chen, L.; Fan, Z.; Chang, J.; Yang, R.; Hou, H.; Guo, H.; Zhang, Y.; Yang, T.; Zhou, C.; Sui, Q.; Chen, Z.; Zheng, C.; Hao, X.; Zhang, K.; Cui, R.; Zhang, Z.; Ma, H.; Ding, Y.; Zhang, N.; Lu, X.; Luo, X.; Jiang, H.; Zhang, S.; Zheng, M. Sequence-Based Drug Design as a Concept in Computational Drug Design. Nat. Commun. 2023, 14 (1), 4217. 10.1038/s41467-023-39856-w.

(15) Ahmed, K. T.; Ansari, M. I.; Zhang, W. DTI-LM: Language Model Powered Drug–Target Interaction Prediction. Bioinformatics 2024, 40 (9), btae533. 10.1093/bioinformatics/btae533.

(16) Jumper, J.; Evans, R.; Pritzel, A.; Green, T.; Figurnov, M.; Ronneberger, O.; Tunyasuvunakool, K.; Bates, R.; Žídek, A.; Potapenko, A.; Bridgland, A.; Meyer, C.; Kohl, S. A. A.; Ballard, A. J.; Cowie, A.; Romera-Paredes, B.; Nikolov, S.; Jain, R.; Adler, J.; Back, T.; Petersen, S.; Reiman, D.; Clancy, E.; Zielinski, M.; Steinegger, M.; Pacholska, M.; Berghammer, T.; Bodenstein, S.; Silver, D.; Vinyals, O.; Senior, A. W.; Kavukcuoglu, K.; Kohli, P.; Hassabis, D. Highly Accurate Protein Structure Prediction with AlphaFold. Nature 2021, 596 (7873), 583–589. 10.1038/s41586-021-03819-2.

(17) Abramson, J.; Adler, J.; Dunger, J.; Evans, R.; Green, T.; Pritzel, A.; Ronneberger, O.; Willmore, L.; Ballard, A. J.; Bambrick, J.; Bodenstein, S. W.; Evans, D. A.; Hung, C.-C.; O’Neill, M.; Reiman, D.; Tunyasuvunakool, K.; Wu, Z.; Žemgulytė, A.; Arvaniti, E.; Beattie, C.; Bertolli, O.; Bridgland, A.; Cherepanov, A.; Congreve, M.; Cowen-Rivers, A. I.; Cowie, A.; Figurnov, M.; Fuchs, F. B.; Gladman, H.; Jain, R.; Khan, Y. A.; Low, C. M. R.; Perlin, K.; Potapenko, A.; Savy, P.; Singh, S.; Stecula, A.; Thillaisundaram, A.; Tong, C.; Yakneen, S.; Zhong, E. D.; Zielinski, M.; Žídek, A.; Bapst, V.; Kohli, P.; Jaderberg, M.; Hassabis, D.; Jumper, J. M. Accurate Structure Prediction of Biomolecular Interactions with AlphaFold 3. Nature 2024, 630 (8016), 493–500. 10.1038/s41586-024-07487-w.

(18) Passaro, S.; Corso, G.; Wohlwend, J.; Reveiz, M.; Thaler, S.; Somnath, V. R.; Getz, N.; Portnoi, T.; Roy, J.; Stark, H.; Kwabi-Addo, D.; Beaini, D.; Jaakkola, T.; Barzilay, R. Boltz-2: Towards Accurate and Efficient Binding Affinity Prediction. bioRxiv 2025, 2025.06.14.659707. 10.1101/2025.06.14.659707.

(19) Satorras, V. G.; Hoogeboom, E.; Welling, M. E(n) Equivariant Graph Neural Networks. In Proceedings of the 38th International Conference on Machine Learning; PMLR, 2021; pp 9323–9332.

(20) Wallach, I.; Dzamba, M.; Heifets, A. AtomNet: A Deep Convolutional Neural Network for Bioactivity Prediction in Structure-Based Drug Discovery. arXiv October 10, 2015. 10.48550/arXiv.1510.02855.

(21) Atz, K.; Grisoni, F.; Schneider, G. Geometric Deep Learning on Molecular Representations. *Nat*. Mach. Intell. 2021, 3 (12), 1023–1032. 10.1038/s42256-021-00418-8.

(22) Ramsundar, B.; Liu, B.; Wu, Z.; Verras, A.; Tudor, M.; Sheridan, R. P.; Pande, V. Is Multitask Deep Learning Practical for Pharma? J. Chem. Inf. Model. 2017, 57 (8), 2068–2076. 10.1021/acs.jcim.7b00146.

(23) Liu, T.; Lin, Y.; Wen, X.; Jorissen, R. N.; Gilson, M. K. BindingDB: A Web-Accessible Database of Experimentally Determined Protein–Ligand Binding Affinities. Nucleic Acids Res. 2007, 35 (Database issue), D198– D201. 10.1093/nar/gkl999.

(24) van Kempen, M.; Kim, S. S.; Tumescheit, C.; Mirdita, M.; Lee, J.; Gilchrist, C. L. M.; Söding, J.; Steinegger, M. Fast and Accurate Protein Structure Search with Foldseek. Nat. Biotechnol. 2024, 42 (2), 243–246. 10.1038/s41587-023-01773-0.

(25) Arevalo, J.; Solorio, T.; Montes-y-Gómez, M.; González, F. A. Gated Multimodal Units for Information Fusion. arXiv February 7, 2017. 10.48550/arXiv.1702.01992.

(26) Weininger, D. SMILES, a Chemical Language and Information System. 1. Introduction to Methodology and Encoding Rules. J. Chem. Inf. Comput. Sci. 1988, 28 (1), 31–36. 10.1021/ci00057a005.

(27) Kipf, T. N.; Welling, M. Semi-Supervised Classification with Graph Convolutional Networks. arXiv February 22, 2017. 10.48550/arXiv.1609.02907.

(28) Heinzinger, M.; Weissenow, K.; Sanchez, J. G.; Henkel, A.; Mirdita, M.; Steinegger, M.; Rost, B. Bilingual Language Model for Protein Sequence and Structure. NAR Genomics Bioinforma. 2024, 6 (4), lqae150. 10.1093/nargab/lqae150.

(29) Vaswani, A.; Shazeer, N.; Parmar, N.; Uszkoreit, J.; Jones, L.; Gomez, A. N.; Kaiser, L.; Polosukhin, I. Attention Is All You Need. arXiv August 2, 2023. 10.48550/arXiv.1706.03762.

(30) Wallach, I.; Heifets, A. Most Ligand-Based Classification Benchmarks Reward Memorization Rather than Generalization. J. Chem. Inf. Model. 2018, 58 (5), 916–932. 10.1021/acs.jcim.7b00403.

(31) Corbin, J. D.; Francis, S. H. Cyclic GMP Phosphodiesterase-5: Target of Sildenafil*. J. Biol. Chem. 1999, 274 (20), 13729–13732. 10.1074/jbc.274.20.13729.

(32) Stavrou, E. X.; Fang, C.; Merkulova, A.; Alhalabi, O.; Grobe, N.; Antoniak, S.; Mackman, N.; Schmaier, A. H. Reduced Thrombosis in Klkb1−/− Mice Is Mediated by Increased Mas Receptor, Prostacyclin, Sirt1, and KLF4 and Decreased Tissue Factor. Blood 2015, 125 (4), 710–719. 10.1182/blood-2014-01-550285.

(33) Houtkooper, R. H.; Pirinen, E.; Auwerx, J. Sirtuins as Regulators of Metabolism and Healthspan. Nat. Rev. Mol. Cell Biol. 2012, 13 (4), 225–238. 10.1038/nrm3293.

(34) Kotian, P. L.; Wu, M.; Vadlakonda, S.; Chintareddy, V.; Lu, P.; Juarez, L.; Kellogg-Yelder, D.; Chen, X.; Muppa, S.; Chambers-Wilson, R.; Davis Parker, C.; Williams, J.; Polach, K. J.; Zhang, W.; Raman, K.; Babu, Y. S. Berotralstat (BCX7353): Structure-Guided Design of a Potent, Selective, and Oral Plasma Kallikrein Inhibitor to Prevent Attacks of Hereditary Angioedema (HAE). J. Med. Chem. 2021, 64 (17), 12453–12468. 10.1021/acs.jmedchem.1c00511.

(35) MOE 2024.06 and 2024.0601 Release Notes. https://www.chemcomp.com/release_notes/moe20240601/rnotes.htm (accessed 2026-02-03).

(36) Kotera, J.; Mochida, H.; Inoue, H.; Noto, T.; Fujishige, K.; Sasaki, T.; Kobayashi, T.; Kojima, K.; Yee, S.; Yamada, Y.; Kikkawa, K.; Omori, K. Avanafil, a Potent and Highly Selective Phosphodiesterase-5 Inhibitor for Erectile Dysfunction. J. Urol. 2012, 188 (2), 668–674. 10.1016/j.juro.2012.03.115.

(37) Ng, T. I.; Tripathi, R.; Reisch, T.; Lu, L.; Middleton, T.; Hopkins, T. A.; Pithawalla, R.; Irvin, M.; Dekhtyar, T.; Krishnan, P.; Schnell, G.; Beyer, J.; McDaniel, K. F.; Ma, J.; Wang, G.; Jiang, L.-J.; Or, Y. S.; Kempf, D.; Pilot-Matias, T.; Collins, C. In Vitro Antiviral Activity and Resistance Profile of the Next-Generation Hepatitis C Virus NS3/4A Protease Inhibitor Glecaprevir. Antimicrob. Agents Chemother. 2018, 62 (1), e01620–17. 10.1128/AAC.01620-17.

(38) Onrust, S. V.; Lamb, H. M. Valrubicin. Drugs Aging 1999, 15 (1), 69–75. 10.2165/00002512-199915010-00006.

(39) Wang, H.; Ye, M.; Robinson, H.; Francis, S. H.; Ke, H. Conformational Variations of Both Phosphodiesterase-5 and Inhibitors Provide the Structural Basis for the Physiological Effects of Vardenafil and Sildenafil. Mol. Pharmacol. 2008, 73 (1), 104–110. 10.1124/mol.107.040212.

(40) Tang, J.; Yu, C. L.; Williams, S. R.; Springman, E.; Jeffery, D.; Sprengeler, P. A.; Estevez, A.; Sampang, J.; Shrader, W.; Spencer, J.; Young, W.; McGrath, M.; Katz, B. A. Expression, Crystallization, and Three-Dimensional Structure of the Catalytic Domain of Human Plasma Kallikrein. J. Biol. Chem. 2005, 280 (49), 41077–41089. 10.1074/jbc.M506766200.

(41) Li, M.; Srp, J.; Gustchina, A.; Dauter, Z.; Mares, M.; Wlodawer, A. Crystal Structures of the Complex of a Kallikrein Inhibitor from Bauhinia Bauhinioides with Trypsin and Modeling of Kallikrein Complexes. Acta Crystallogr. Sect. Struct. Biol. 2019, 75 (Pt 1), 56–69. 10.1107/S2059798318016492.

(42) Jin, L.; Wei, W.; Jiang, Y.; Peng, H.; Cai, J.; Mao, C.; Dai, H.; Choy, W.; Bemis, J. E.; Jirousek, M. R.; Milne, J. C.; Westphal, C. H.; Perni, R. B. Crystal Structures of Human SIRT3 Displaying Substrate-Induced Conformational Changes. J. Biol. Chem. 2009, 284 (36), 24394–24405. 10.1074/jbc.M109.014928.

(43) Sung, B.-J.; Hwang, K. Y.; Jeon, Y. H.; Lee, J. I.; Heo, Y.-S.; Kim, J. H.; Moon, J.; Yoon, J. M.; Hyun, Y.-L.; Kim, E.; Eum, S. J.; Park, S.-Y.; Lee, J.-O.; Lee, T. G.; Ro, S.; Cho, J. M. Structure of the Catalytic Domain of Human Phosphodiesterase 5 with Bound Drug Molecules. Nature 2003, 425 (6953), 98–102. 10.1038/nature01914.

(44) Druker, B. J.; Tamura, S.; Buchdunger, E.; Ohno, S.; Segal, G. M.; Fanning, S.; Zimmermann, J.; Lydon, N. B. Effects of a Selective Inhibitor of the Abl Tyrosine Kinase on the Growth of Bcr-Abl Positive Cells. Nat. Med. 1996, 2 (5), 561–566. 10.1038/nm0596-561.

(45) Vig, J. A Multiscale Visualization of Attention in the Transformer Model. arXiv June 12, 2019. 10.48550/arXiv.1906.05714.

(46) Gorre, M. E.; Mohammed, M.; Ellwood, K.; Hsu, N.; Paquette, R.; Rao, P. N.; Sawyers, C. L. Clinical Resistance to STI-571 Cancer Therapy Caused by BCR-ABL Gene Mutation or Amplification. Science 2001, 293 (5531), 876–880. 10.1126/science.1062538.

(47) Schindler, T.; Bornmann, W.; Pellicena, P.; Miller, W. T.; Clarkson, B.; Kuriyan, J. Structural Mechanism for STI-571 Inhibition of Abelson Tyrosine Kinase. Science 2000, 289 (5486), 1938–1942. 10.1126/science.289.5486.1938.

(48) Welch, B. L. The Generalization of ‘Student’s’ Problem When Several Different Population Variances Are Involved. Biometrika 1947, 34 (1/2), 28–35. 10.2307/2332510.

(49) Branford, S.; Rudzki, Z.; Walsh, S.; Parkinson, I.; Grigg, A.; Szer, J.; Taylor, K.; Herrmann, R.; Seymour, J. F.; Arthur, C.; Joske, D.; Lynch, K.; Hughes, T. Detection of BCR-ABL Mutations in Patients with CML Treated with Imatinib Is Virtually Always Accompanied by Clinical Resistance, and Mutations in the ATP Phosphate-Binding Loop (P-Loop) Are Associated with a Poor Prognosis. Blood 2003, 102 (1), 276–283. 10.1182/blood-2002-09-2896.

(50) Ohren, J. F.; Chen, H.; Pavlovsky, A.; Whitehead, C.; Zhang, E.; Kuffa, P.; Yan, C.; McConnell, P.; Spessard, C.; Banotai, C.; Mueller, W. T.; Delaney, A.; Omer, C.; Sebolt-Leopold, J.; Dudley, D. T.; Leung, I. K.; Flamme, C.; Warmus, J.; Kaufman, M.; Barrett, S.; Tecle, H.; Hasemann, C. A. Structures of Human MAP Kinase Kinase 1 (MEK1) and MEK2 Describe Novel Noncompetitive Kinase Inhibition. Nat. Struct. Mol. Biol. 2004, 11 (12), 1192–1197. 10.1038/nsmb859.

(51) The UniProt Consortium. UniProt: The Universal Protein Knowledgebase in 2023. Nucleic Acids Res. 2023, 51 (D1), D523–D531. 10.1093/nar/gkac1052.

(52) Kane, J.; Honigfeld, G.; Singer, J.; Meltzer, H. Clozapine for the Treatment-Resistant Schizophrenic. A Double-Blind Comparison with Chlorpromazine. Arch. Gen. Psychiatry 1988, 45 (9), 789–796. 10.1001/archpsyc.1988.01800330013001.

(53) McTavish, D.; Benfield, P. Clomipramine. An Overview of Its Pharmacological Properties and a Review of Its Therapeutic Use in Obsessive Compulsive Disorder and Panic Disorder. Drugs 1990, 39 (1), 136–153. 10.2165/00003495-199039010-00010.

(54) Meltzer, H. Y. Clinical Studies on the Mechanism of Action of Clozapine: The Dopamine-Serotonin Hypothesis of Schizophrenia. Psychopharmacology (Berl.) 1989, 99 *Suppl*, S18–27. 10.1007/BF00442554.

(55) Tatsumi, M.; Groshan, K.; Blakely, R. D.; Richelson, E. Pharmacological Profile of Antidepressants and Related Compounds at Human Monoamine Transporters. Eur. J. Pharmacol. 1997, 340 (2–3), 249–258. 10.1016/s0014-2999(97)01393-9.

(56) Richelson, E. Receptor Pharmacology of Neuroleptics: Relation to Clinical Effects. J. Clin. Psychiatry 1999, 60 *Suppl 10*, 5–14.

(57) Gillman, P. K. Tricyclic Antidepressant Pharmacology and Therapeutic Drug Interactions Updated. Br. J. Pharmacol. 2007, 151 (6), 737–748. 10.1038/sj.bjp.0707253.

(58) Gentile, F.; Agrawal, V.; Hsing, M.; Ton, A.-T.; Ban, F.; Norinder, U.; Gleave, M. E.; Cherkasov, A. Deep Docking: A Deep Learning Platform for Augmentation of Structure Based Drug Discovery. ACS Cent. Sci. 2020, 6 (6), 939–949. 10.1021/acscentsci.0c00229.

(59) Ultra-large library docking for discovering new chemotypes | Nature. https://www.nature.com/articles/s41586-019-0917-9 (accessed 2026-03-12).

(60) Buel, G. R.; Walters, K. J. Can AlphaFold2 Predict the Impact of Missense Mutations on Structure? Nat. Struct. Mol. Biol. 2022, 29 (1), 1–2. 10.1038/s41594-021-00714-2.

(61) Karniadakis, G. E.; Kevrekidis, I. G.; Lu, L.; Perdikaris, P.; Wang, S.; Yang, L. Physics-Informed Machine Learning. Nat. Rev. Phys. 2021, 3 (6), 422–440. 10.1038/s42254-021-00314-5.

(62) Olivecrona, M.; Blaschke, T.; Engkvist, O.; Chen, H. Molecular De-Novo Design through Deep Reinforcement Learning. J. Cheminformatics 2017, 9 (1), 48. 10.1186/s13321-017-0235-x.

(63) Huang, K.; Fu, T.; Glass, L. M.; Zitnik, M.; Xiao, C.; Sun, J. DeepPurpose: A Deep Learning Library for Drug–Target Interaction Prediction. Bioinformatics 2021, 36 (22–23), 5545–5547. 10.1093/bioinformatics/btaa1005.

(64) Landrum, G.; others. RDKit: A Software Suite for Cheminformatics, Computational Chemistry, and Predictive Modeling. Greg Landrum 2013, 8 (31.10), 5281.

(65) Hendrycks, D.; Gimpel, K. Gaussian Error Linear Units (GELUs). arXiv June 6, 2023. 10.48550/arXiv.1606.08415.

(66) Ba, J. L.; Kiros, J. R.; Hinton, G. E. Layer Normalization. arXiv July 21, 2016. 10.48550/arXiv.1607.06450.

(67) Paszke, A.; Gross, S.; Massa, F.; Lerer, A.; Bradbury, J.; Chanan, G.; Killeen, T.; Lin, Z.; Gimelshein, N.; Antiga, L.; Desmaison, A.; Kopf, A.; Yang, E.; DeVito, Z.; Raison, M.; Tejani, A.; Chilamkurthy, S.; Steiner, B.; Fang, L.; Bai, J.; Chintala, S. PyTorch: An Imperative Style, High-Performance Deep Learning Library. In Advances in Neural Information Processing Systems; Curran Associates, Inc., 2019; Vol. 32.

(68) Bergstra, J.; Bengio, Y. Random Search for Hyper-Parameter Optimization. J Mach Learn Res 2012, 13 (null), 281–305.

(69) Srivastava, N.; Hinton, G.; Krizhevsky, A.; Sutskever, I.; Salakhutdinov, R. Dropout: A Simple Way to Prevent Neural Networks from Overfitting. J. Mach. Learn. Res. 2014, 15 (56), 1929–1958.

(70) Liu, L.; Jiang, H.; He, P.; Chen, W.; Liu, X.; Gao, J.; Han, J. On the Variance of the Adaptive Learning Rate and Beyond. arXiv October 26, 2021. 10.48550/arXiv.1908.03265.

(71) Sergeev, A.; Balso, M. D. Horovod: Fast and Easy Distributed Deep Learning in TensorFlow. arXiv February 21, 2018. 10.48550/arXiv.1802.05799.

(72) Steck, H.; Krishnapuram, B.; Dehing-oberije, C.; Lambin, P.; Raykar, V. C. On Ranking in Survival Analysis: Bounds on the Concordance Index. In Advances in Neural Information Processing Systems; Curran Associates, Inc., 2007; Vol. 20.

(73) Davis, J.; Goadrich, M. The Relationship between Precision-Recall and ROC Curves. In Proceedings of the 23rd international conference on Machine learning; ICML ’06; Association for Computing Machinery: New York, NY, USA, 2006; pp 233–240. 10.1145/1143844.1143874.

(74) Wilcoxon, F. Individual Comparisons by Ranking Methods. Biom. Bull. 1945, 1 (6), 80–83. 10.2307/3001968.

(75) Zhang, J.; Krishnan, R.; Arnold, C. S.; Mattsson, E.; Kilpatrick, J. M.; Bantia, S.; Dehghani, A.; Boudreaux, B.; Gupta, S. N.; Kotian, P. L.; Chand, P.; Babu, Y. S. Discovery of Highly Potent Small Molecule Kallikrein Inhibitors. Med. Chem. Shariqah United Arab Emir. 2006, 2 (6), 545–553. 10.2174/1573406410602060545.

(76) He, W.; Newman, J. C.; Wang, M. Z.; Ho, L.; Et., A. Mitochondrial Sirtuins: Regulators of Protein Acylation and Metabolism. Trends Endocrinol. Metab. 2012. 10.1016/j.tem.2012.07.004.

(77) Naïm, M.; Bhat, S.; Rankin, K. N.; Dennis, S.; Chowdhury, S. F.; Siddiqi, I.; Drabik, P.; Sulea, T.; Bayly, C. I.; Jakalian, A.; Purisima, E. O. Solvated Interaction Energy (SIE) for Scoring Protein−Ligand Binding Affinities. 1. Exploring the Parameter Space. J. Chem. Inf. Model. 2007, 47 (1), 122–133. 10.1021/ci600406v.

(78) Berman, H. M.; Westbrook, J.; Feng, Z.; Gilliland, G.; Bhat, T. N.; Weissig, H.; Shindyalov, I. N.; Bourne, P. E. The Protein Data Bank. Nucleic Acids Res. 2000, 28 (1), 235–242. 10.1093/nar/28.1.235.

(79) Mendez, D.; Gaulton, A.; Bento, A. P.; Chambers, J.; De Veij, M.; Félix, E.; Magariños, M. P.; Mosquera, J. F.; Mutowo, P.; Nowotka, M.; Gordillo-Marañón, M.; Hunter, F.; Junco, L.; Mugumbate, G.; Rodriguez-Lopez, M.; Atkinson, F.; Bosc, N.; Radoux, C. J.; Segura-Cabrera, A.; Hersey, A.; Leach, A. R. ChEMBL: Towards Direct Deposition of Bioassay Data. Nucleic Acids Res. 2019, 47 (D1), D930–D940. 10.1093/nar/gky1075.

(80) Jabbour, E.; Kantarjian, H. Chronic Myeloid Leukemia: 2022 Update on Diagnosis, Therapy, and Monitoring. Am. J. Hematol. 2022, 97 (9), 1236–1256. 10.1002/ajh.26642.

(81) Kim, S.; Chen, J.; Cheng, T.; Gindulyte, A.; He, J.; He, S.; Li, Q.; Shoemaker, B. A.; Thiessen, P. A.; Yu, B.; Zaslavsky, L.; Zhang, J.; Bolton, E. E. PubChem 2023 Update. Nucleic Acids Res. 2023, 51 (D1), D1373– D1380. 10.1093/nar/gkac956.

(82) Pettersen, E. F.; Goddard, T. D.; Huang, C. C.; Meng, E. C.; Couch, G. S.; Croll, T. I.; Morris, J. H.; Ferrin, T. E. UCSF ChimeraX: Structure Visualization for Researchers, Educators, and Developers. Protein Sci. Publ. Protein Soc. 2021, 30 (1), 70–82. 10.1002/pro.3943.

