## Supplementary Tables S1-S4, Supplementary Figure S1-S3 for "Pan–Pharmacological Drug–Target Interaction Prediction with 3D–Informed Protein Encoding at Scale"

**Supplementary Table S1: Statistics of the BindingDB datasets used for model development and evaluation.**

| Dataset | $K_i$ | $IC_{50}$ | $EC_{50}$ | $K_d$ | Total (unique pairs) |
|---|---|---|---|---|---|
| **BindingDB 2024 (Full)** | 564,724 | 1,834,763 | 258,957 | 108,467 | 2,761,182 |
| **Training set (BindingDB 2023, Seed 42)** | 370,636 | 1,129,488 | 166,290 | 63,106 | 1,725,174 |
| **Validation set (BindingDB 2023, Seed 42)** | 66,579 | 200,514 | 29,927 | 10,795 | 306,996 |
| **Label reversal test set** | 66,560 | 142,974 | 21,908 | 19,946 | 250,827 |

| | | | | | |
|---|---|---|---|---|---|
| **(BindingDB 2023)** | | | | | |
| **Temporal validation set (BindingDB 2024)** | 46,943 | 325,129 | 34,876 | 16,852 | 423,647 |

**Supplementary Table S2: Atom–level chemical features utilized for graph–based compound representation**

| Feature category | Dimensions | Description / Encoding details |
|---|---|---|
| Atom symbol | 10 | One–of–K encoding: C, N, O, F, P, S, Cl, Br, I, or others |
| Degree | 8 | One–of–K encoding of the number of directly bonded neighbors: 0, 1, 2, 3, 4, 5, 6, or others |
| Formal charge | 1 | Integer value (e.g., –1, 0, 1) |

| | | |
|---|---|---|
| Radical electrons | 1 | Integer value |
| Hybridization type | 6 | One–of–K encoding: SP, SP2, SP3, SP3D, SP3D2, or others |
| Aromaticity | 1 | 0 (non-aromatic) or 1 (aromatic) |
| Total number of Hs | 5 | One–of–K encoding of hydrogen atoms (explicit and implicit): 0, 1, 2, 3, or 4 |
| Chirality | 3 | One–of–K encoding of CIP codes (R or S) and a binary indicator for potential chirality |
| Total | 35 | |

**Supplementary Table S3: Hyperparameter search space and selected configurations for the OmniBind architecture**

| Hyperparameter | Search range | Selected value |
|---|---|---|
| Number of sequence encoder layers | 1, 2, 3 | 2 |
| Number of structure encoder layers | 1, 2, 3 | 2 |

| | | |
|---|---|---|
| Number of decoder layers | 1, 2, 3, 4, 5, 6 | 5 |
| Number of attention heads | 4, 8, 12, 16 | 8 |
| Hidden dimension | 128, 256, 512, 768 | 256 |
| Dropout rate | 0.1, 0.2, 0.3 | 0.1 |

**Supplementary Table S4: Performance of protein input representations across individual pharmacological metrics**

RMSE is reported for each of the four pharmacological endpoints ($pK_i, pIC_{50}, pEC_{50}$, and $pK_d$). The "Avg (3 labels)" represents the mean RMSE of $pK_i, pIC_{50}$, and $pEC_{50}$; "Avg (4 labels)" additionally includes $pK_d$. $pK_d$ was excluded from the primary average reported in Table 1 due to its substantially smaller sample size and higher inter-replicate variance in BindingDB. Values represent the mean ± standard deviation across five independent runs.

| **Protein input representation** | **RMSE ($pK_i$)** | **RMSE ($pIC_{50}$)** | **RMSE ($pEC_{50}$)** | **RMSE ($pK_d$)** | **Avg (3 labels)** | **Avg (4 labels)** |
|---|---|---|---|---|---|---|

| | | | | | | |
|---|---|---|---|---|---|---|
| Only 3Di | 1.051 ± 0.017 | 1.026 ± 0.012 | 1.045 ± 0.022 | 1.837 ± 0.122 | 1.041 | 1.240 |
| Only amino acids | 1.063 ± 0.011 | 1.026 ± 0.010 | 1.059 ± 0.028 | 1.858 ± 0.097 | 1.049 | 1.252 |
| 3Di + amino acids (add) | 1.048 ± 0.009 | **1.024 ± 0.011** | 1.053 ± 0.031 | **1.763 ± 0.056** | 1.042 | **1.222** |
| 3Di + amino acids (gate fusion) | **1.045 ± 0.010** | 1.031 ± 0.012 | **1.035 ± 0.017** | 1.830 ± 0.103 | **1.037** | 1.235 |

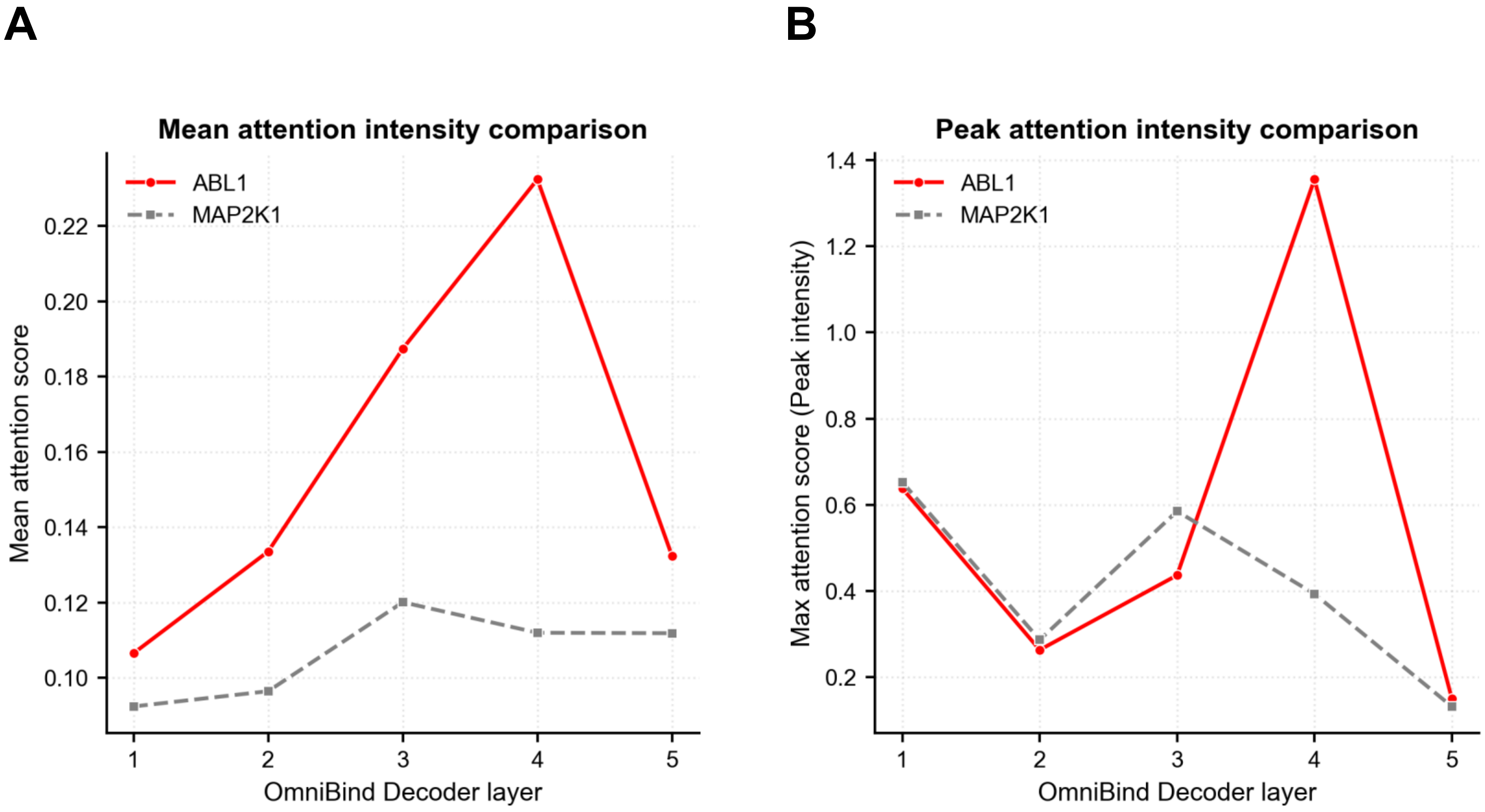


**Kawaharada et al., Supplementary Figure S1**

**Supplementary Figure S1: Negative control attention analysis for the non–binding MAP2K1–imatinib pair.**

**(A)** Mean cross-attention scores across the five OmniBind decoder layers for the ABL1–imatinib (red) and MAP2K1–imatinib (gray) pairs. While ABL1–imatinib attention scores increase progressively and peak in layer 4, MAP2K1–imatinib scores remain consistently low across all layers, with no layer–specific signal concentration. **(B)** Peak cross-attention scores across decoder layers for the same two pairs. The ABL1–imatinib pair shows a pronounced spike in layer 4, whereas MAP2K1–imatinib exhibits no comparable localized signal. Together, these results confirm that the elevated and layer–specific attention patterns observed for ABL1–imatinib reflect genuine binding recognition rather than a non–specific property of the decoder architecture.

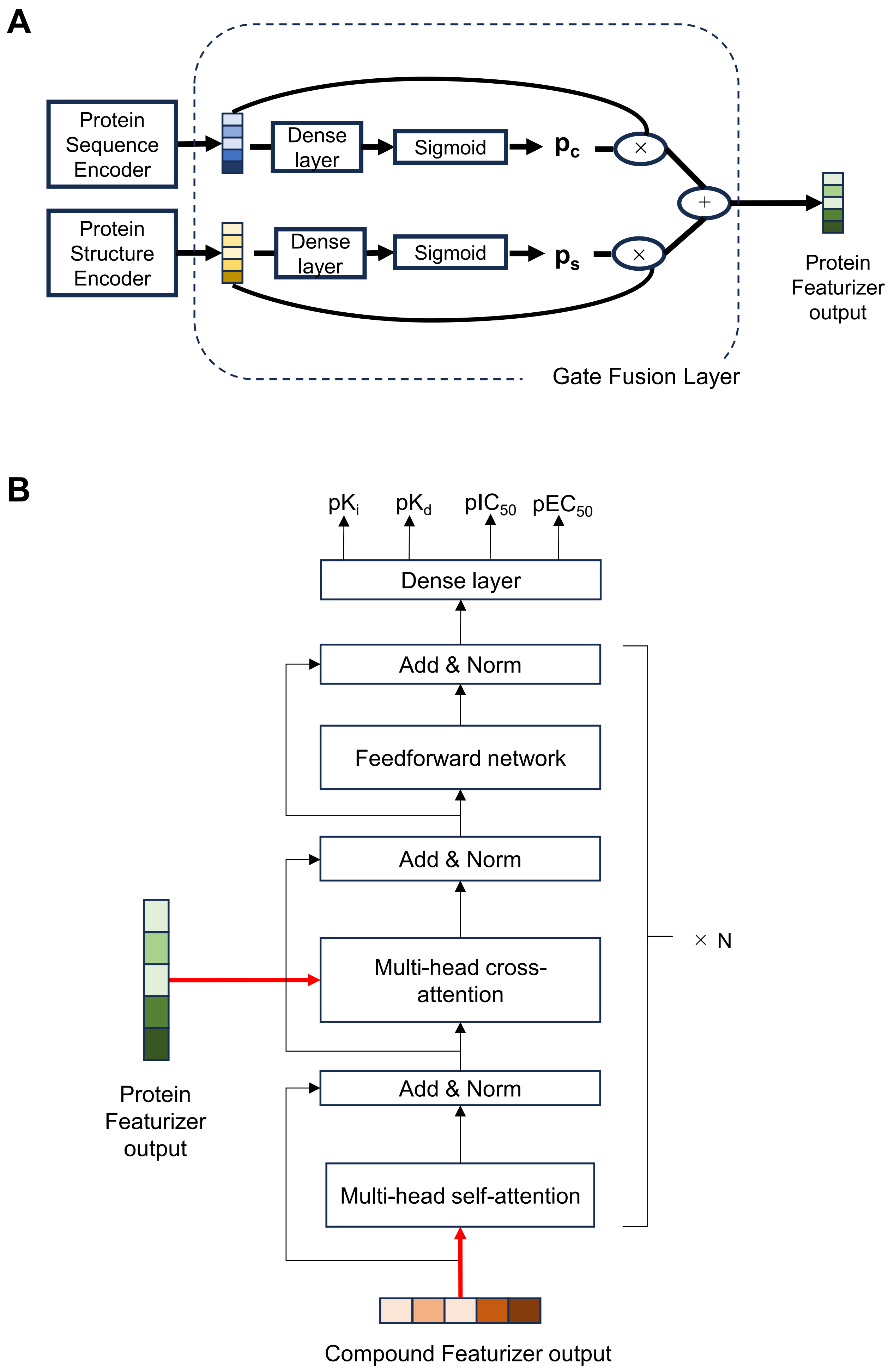


**Kawaharada et al., Supplementary Figure S2**

**Supplementary Figure S2: Architectural details of the gate fusion layer and OmniBind decoder**

**(A)** Schematic of the gate fusion layer. Representations from the Protein Sequence and Protein Structure Encoders are each passed through a dense layer followed by sigmoid activation to produce gating coefficients denoted as $p_c$ and $p_s$. These coefficients are applied as element–wise weights to the respective encoder outputs, and the weighted representations are summed to produce the fused protein featurizer output (The mathematical formulation is provided in the Protein Featurizer section of Materials and Methods). **(B)** Schematic of the OmniBind decoder block. Multi–head self–attention is first applied to the compound embeddings, followed by Add & Norm. The decoder then performs multi–head cross–attention using the fused protein representation as the key and value, followed by Add & Norm and a feedforward network, enabling the model to learn compound–protein interactions. The resulting features are passed through a fully connected layer to simultaneously output four pharmacological metrics ($pK_i, pK_d, pIC_{50}$, and $pEC_{50}$).

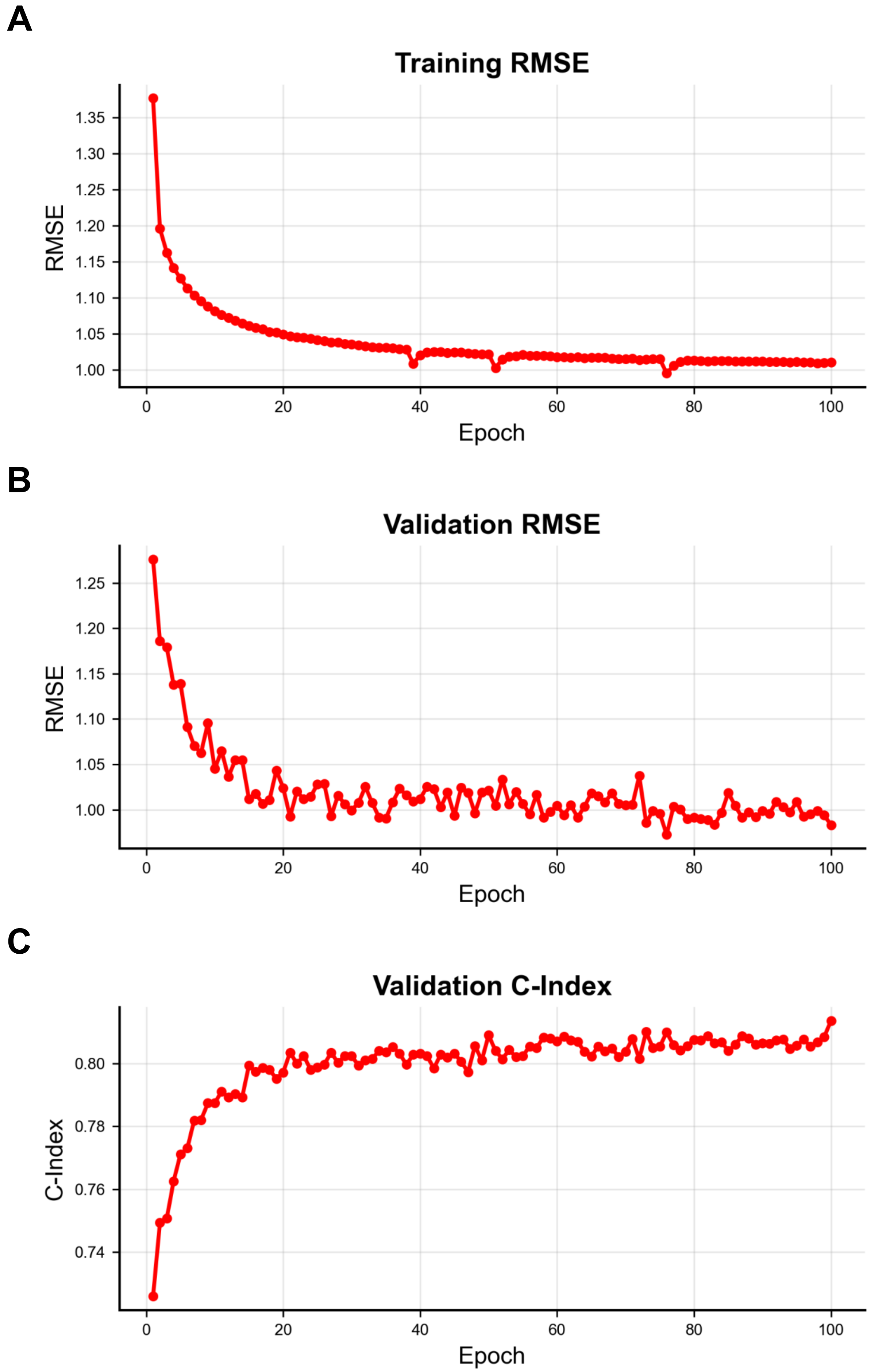


**Kawaharada et al., Supplementary Figure S3**

**Supplementary Figure S3: Training convergence of OmniBind over 100 epochs**

**(A)** Training RMSE as a function of epoch. The loss decreases rapidly in early epochs and stabilizes below 1.05 by epoch 40, indicating stable convergence. **(B)** Validation RMSE as a function of epoch. The error similarly converges to approximately 1.0 without divergence, confirming that the model generalizes to held–out data without overfitting. **(C)** Validation C–Index as a function of epoch. The C–index increases rapidly in early training and plateaus above 0.80, indicating that the model consistently ranks compound–protein pairs by affinity with high accuracy.
